# The pleiotropic effects of the MNK1/2-eIF4E axis support immune suppression and metastasis in a model of postpartum breast cancer

**DOI:** 10.1101/2020.09.07.286815

**Authors:** Qianyu Guo, Margarita Bartish, Christophe Goncalves, Fan Huang, Sai Sakktee Krisna, Samuel E. J. Preston, Audrey Emond, Vivian Zihui Li, Claudia U. Duerr, Yirui Gui, Aurélie Cleret-Buhot, Paméla Thébault, Hanne Lefrère, Liesbeth Lenaerts, Dany Plourde, Jie Su, Barbara C. Mindt, Shannon Hewgill, Tiziana Cotechini, Charles Hindmarch, William Yang, Elie Khoury, Yao Zhan, Valeria Narykina, Yuhong Wei, Giuseppe Floris, Mark Basik, Frédéric Amant, Daniela F. Quail, Réjean Lapointe, Jörg H. Fritz, Sonia V. del Rincón, Wilson H. Miller

## Abstract

**Purpose:** Breast cancer diagnosed within 10 years following childbirth is defined as postpartum breast cancer (PPBC) and is highly metastatic. Interactions between immune cells and other stromal cells within the involuting mammary gland are fundamental in facilitating an aggressive tumor phenotype. The MNK1/2-eIF4E axis promotes the translation of pro-metastatic mRNAs in tumor cells, but its role in modulating the function of non-tumor cells in the PPBC microenvironment, and in particular its activity in human PPBC, has not been explored.

**Experimental design:** We used a combination of in vivo PPBC models and in vitro assays to study the effects of phospho-eIF4E deficiency on the pro-tumor function of select cells of the TME. Furthermore, we employed Imaging Mass Cytometry on PPBC and non-PPBC patient samples, to chart the expression of the MNK1/2-eIF4E axis components in the TME.

**Results:** Here, we show that phospho-eIF4E deficient (eIF4E^S209A^) PPBC mice are protected against lung metastasis and reveal differences in the lung immune microenvironment of the WT and eIF4E^S209A^ mice. Moreover, we show that the expression of fibroblast-derived IL-33, an alarmin known to induce invasion, was repressed upon MNK1/2-eIF4E axis inhibition. Imaging Mass Cytometry results indicated that human PPBC contain phospho-eIF4E high-expressing tumor cells and CD8+ T cells displaying an activated dysfunctional phenotype. Finally, we block lung metastasis in PPBC mice, using combined MNK1/2 inhibition and anti-PD-1 therapy.

**Conclusions:** These findings implicate the involvement of the MNK1/2-eIF4E axis during PPBC metastasis and suggest a promising immunomodulatory route to enhance the efficacy of immunotherapy by blocking phospho-eIF4E.

**Translational relevance:** Postpartum breast cancer (PPBC) is highly aggressive. It is hypothesized that involution-induced changes in the postpartum breast microenvironment, which include an influx of inflammatory immune cells and activation of resident fibroblasts, facilitate the invasiveness of an existing neoplasm. We used imaging mass cytometry to do an in-depth profiling of the MNK1-eIF4E axis in the TME of a unique cohort of PPBC and non-PPBC patients. We observed patterns of phospho-eIF4E in non-tumor cells that were specific to the TME of PPBC. We also noted that the CD8^+^ T cells present in PPBC express an activated dysfunctional phenotype characterized by the co-expression of HLA-DR and PD-1. This study represents a first look at the expression of the MNK1-eIF4E axis in the stromal cells of metastatic breast cancer and has therapeutic implications as we show, in an animal model of PPBC, that MNK1/2 inhibition can be used to sensitize tumors to anti-PD1 immunotherapy.

## Introduction

Postpartum breast cancer (PPBC) is defined as breast cancer diagnosed within 10 years of parturition (1). Given its highly metastatic nature (1,2), the patient prognosis is poor. The physiological process of mammary gland (MG) involution, the remodeling of the breast tissue back to its pre-pregnant state, has been hypothesized to cause premalignant epithelial cells to adopt invasive properties (2). Involution, akin to the process of wound healing, is accompanied by an orchestrated immune cell infiltration to the mammary gland (3). Data from murine PPBC models suggest that interactions between innate and adaptive immune cells are fundamental in establishing a suppressed microenvironment that is favorable for metastatic spread. In addition to immune cells, fibroblasts are another critical component of the involuting mammary gland (4). During involution, fibroblasts enter an activated state characterized by an increased expression of genes regulating collagen deposition and the production of immunosuppressive chemokines (4). Activated fibroblasts help to remodel the mammary gland back to a non-lactational state, but in the context of PPBC, they may facilitate evasion from anti-tumor immunity.

Cancer cell invasiveness is regulated by growth factors, cytokines, and chemokines produced by tumor cells and associated stromal cells within the TME. Recently the IL-33/ST2 signaling axis has come to the forefront as an important mediator of metastasis. IL-33 is an alarmin cytokine of the IL-1 family, which is involved in inflammation, tissue homeostasis and tumor progression, and signals via binding to the ST2 receptor (5,6). Although there are many cellular sources of IL-33, its secretion by cancer associated fibroblasts has been shown to promote the epithelial-to-mesenchymal like transition and tumor cell invasion (7,8). IL-33 can also promote tumor progression and immune suppression via activation of immune cells such as CD11b^+^Gr1^+^ myeloid-derived suppressor cells (MDSCs) (9) and innate lymphoid cells type 2 (ILC2) (10). Once activated, these cells serve as potent inhibitors of cytotoxic T cell tumor infiltration and anti-tumor function (11,12).

Regulation of gene expression at the level of mRNA translation initiation is becoming increasingly studied in the field of onco-immunology. Indeed, dysregulation of translational control is a prominent feature of many cancers (13). For example, elevated levels of the eukaryotic translation initiator factor 4E (eIF4E), which binds to the 7-methylguanosine cap at the 5’ end of the mRNA (13,14), are associated with malignancy and poor prognosis in several cancer types (15-20). eIF4E can be phosphorylated at serine 209 (S209) by MAP kinase-interacting serine/threonine-protein kinases 1 and 2 (MNK1/2), and this post-translational modification is essential for its pro-invasive effects (17). Increased MNK1/2 activity has been associated with therapeutic resistance, tumorigenesis, invasion and metastasis (21-27). We and others have previously shown that phosphorylation of eIF4E leads to the translational upregulation of mRNAs, such as *Myc, Mcl1, Mmp3* and *Snai1*, that support tumor cell survival and a pro-invasive phenotype (17,24). Phospho-eIF4E has recently been reported to reinforce the survival of pro-metastatic neutrophils (28). However, there remain large gaps in our understanding of how the regulation of eIF4E phosphorylation impacts the behavior of other immune and non-immune stromal cells found within the breast tumor microenvironment (TME).

Here we show that host phospho-eIF4E has a pleiotropic effect in the TME of an animal model of PPBC, regulating the functions of fibroblasts and ILC2, two cell types critical for the metastatic process. The altered functionality of fibroblasts, in turn, differentially affects tumor cells, to support the immune evasion and metastasis of PPBC tumors. In a pioneering approach, using Imaging Mass Cytometry on a cohort of human PPBC and non-PPBC tumors, we show that the human PPBC TME is characterized by markers of immune dysfunction. Finally, we provide evidence for a potential therapeutic intervention in PPBC, by showing that the combination of the MNK1/2 inhibitor SEL201 and anti-PD1 blockade decreases lung metastasis in a murine model of PPBC.

## Methods

### Mouse Model

Wild-type (WT) BALB/c and C57BL/6N mice were purchased from Charles River Laboratory. eIF4E^S209A/S209A^ BALB/c and eIF4E^S209A/S209A^ C57BL/6N mice were gifts from Dr. Nahum Sonenberg at McGill University and have been previously described (17). PPBC models were set up as previously reported (29,30). Briefly, 6-week old WT or eIF4E^S209A/S209A^ female mice were mated with male mice. Pregnant mice were monitored until new pups were born and allowed to lactate for 11 to 14 days. Pups were removed from the dams causing the dams to undergo forced weaning-induced mammary gland involution. On involution day 1, that is twenty-four hours post-forced weaning, 200,000 66cl4 cells were injected into the inguinal mammary gland of BALB/c mice and tumors were allowed to grow for 14 or 33 days. 200,000 E0771 cells were injected into the mammary gland of C57BL/6N mice for 26 days. For drug treatment experiments, animals were treated at time points indicated in the schematic illustration. SEL201 (Ryvu Therapeutics) was dissolved in DMSO and then diluted in Captisol (Ligand) for administration by oral gavage at 75 mg/kg bodyweight per mouse per day, 5 days per week. The anti-mouse PD-1 monoclonal antibody and IgG isotype control (BioCell) were diluted in PBS and administrated through intraperitoneal injection at 10 mg/kg bodyweight per mouse per day, once per week. Animal experiments were conducted following protocols approved by McGill University Animal Care and Use Committee.

### Cells and Reagents

SEL201 was a generous gift from Dr. Tomasz Rzymski at Ryvu Therapeutics. The 66cl4 and MDA-MB-231 cell lines were kind gifts from Dr. Josie Ursini-Siegel at McGill University. The E0771 cell line was purchased from CH3 BioSystems. 66cl4 was cultured in RPMI with 10% FBS and antibiotics. E0771 was cultured in RPMI supplemented with 10mM HEPES, 10% FBS and antibiotics. WT and eIF4E^S209A/S209A^ (referred to as eIF4E^S209A^) primary mammary gland fibroblasts were obtained by digesting minced mammary glands pooled from 3-4 donor mice in 1 mg/ml Collagenase IV in DMEM Advanced F12 for 1h at 37 °C, passing the suspension through a 70 µm cell strainer and centrifuging at 300g for 10 minutes. The pelleted cell suspension including fibroblasts was plated in DMEM supplemented with 10% FBS and antibiotics (1x pen/strep, wisent). To enrich for fibroblasts, culture medium was changed 30 minutes after plating. Cells were expanded for 8-9 days. Conditioned media was prepared by thoroughly washing away culture media and culturing the fibroblasts in serum-free DMEM Advanced F12 for 48h. Presence of secreted IL-33 in the conditioned media was visualized on a Proteome Profiler Mouse XL Cytokine Array (R&D Systems). The concentration of IL-33 secreted in the conditioned medium was measured on a V-PLEX Mouse Cytokine 19-Plex Kit (Meso Scale Diagnostics) and normalized to total protein input measured by Nanodrop. WT and eIF4E^S209A/S209A^ (termed eIF4E^S209A^) mouse embryonic fibroblasts (MEFs) have been described previously (17,24). Cancer-associated fibroblasts (CAFs) derived from patient breast cancer were obtained in collaboration with Dr. Mark Basik at McGill University as previously described (31). The collection and use of human tissues was approved by the Institutional Review Board (IRB), JGH (No. 05-006), which is in accordance with the Declaration of Helsinki and the Belmont Report. CAFs and MDA-MB-231 cells were cultured with DMEM supplemented with 10% FBS and antibiotics (1x pen/strep, wisent).

### Migration and Invasion and co-culture assays

66cl4 and E0771 cells were seeded at one (migration and invasion assay) or two (co-culture assay) million cells per 10 cm dish on day 1 in full media, then starved overnight by switching them to serum-free media on day 2. For the co-culture assay, 200,000 WT or eIF4E^S209A^ mouse embryonic fibroblasts were seeded into 12-well companion plates on day 2. On day 3, transwells were coated with Collagen I (20 μg/ml) as previously reported (23). 200,000 (migration and invasion) or 50000 (co-culture assay) tumor cells were seeded into the transwells (Corning) and were allowed to migrate and invade for 16h (migration and invasion) or 48h (co-culture). Migrated cells were fixed with 5% glutaraldehyde (Sigma) and stained with 0.5% crystal violet (Sigma) as previously reported (23). Stained cells were then counted and quantified. WT and eIF4E^S209A^ fibroblasts were harvested for WB or qPCR

For experiments with patient-derived CAFs, MDA-MB-231 cells were seeded at 3 million cells per 10cm dish on day 1 in full media, then switched to serum-free media on day 2 and starved overnight. 50,000 patient-derived CAFs were seeded into 6-well companion plates on day 2. On day 3, transwells were coated with Collagen I (20 μg/ml) as previously reported. 200,000 MDA-MB-231 cells were seeded into the transwells and were allowed to migrate and invade towards CAFs for 48hrs. Migrated cells were fixed, stained and quantified as described above. CAFs were harvested for WB.

### Immunohistochemistry (IHC)

Immunohistochemistry and hematoxylin and eosin (H&E) staining were performed as previously described (23). Briefly, tumor and lung sections were stained for IL-33, phospho-eIF4E and CD8, and counterstained with 20% Harris-modified hematoxylin (Fisher). Antibody information is listed in Supplementary Table 2. Slides were scanned and assessed using Spectrum (Aperio Technologies). All animal and patient IHC samples were quantified by QuPath software.

### Immunofluorescence (IF)

IF staining was performed as previously described (32). Briefly, tissues were stained for p-eIF4E, p-MNK1, IL-33, α-smooth muscle actin (α-SMA), CD45 and CD8, and nucleus were labeled with DAPI. Primary and secondary antibodies were listed in Supplementary Table 2. Slides were scanned with an axioscan Z1 slide scanner microscope (Zeiss) using a 20x/0.75NA objective. Images were analyzed using Zen blue software (Zeiss) and Qupath. A detailed description of analysis procedure can be found in supplemental methods.

### Data acquisition by IMC

The study was approved by the ethics committee and in compliance with institutional review board approval from the 2 participating institutions: UZ/KU Leuven (Belgium) and Jewish General Hospital (Canada). Written informed consent was obtained from all patients (nulliparous breast cancer samples, breast cancer samples from patients diagnosed during pregnancy or diagnosed postpartum) and the study conducted in accordance with the Declaration of Helsinki. Human breast cancer samples were arrayed onto a slide, stained using the panel of antibodies listed in Supplementary Table 4, and processed with the Hyperion Imaging System (Fluidigm) by the Single Cell Imaging and Mass Cytometry Analysis Platform (SCIMAP) of the Goodman Cancer Research Centre, McGill University, according to their guidelines. Areas of dimension 1000 × 1000 µm were acquired for 23 sample cores. The resulting data files were stored in MCD binary format. A detailed description of analysis can be found in supplemental methods.

### Statistical Analysis

Prism software (GraphPad) was used to determine statistical significance of differences. Unpaired Student’s t test, one-way ANOVA or two-way ANOVA is used, as appropriate. P values < 0.05 were considered significant. The details of statistical analysis for each experiment are listed in Supplementary Tables 5-6.

## Results

### Loss of eIF4E phosphorylation in the stroma protects against PPBC lung metastasis

We have previously reported that the absence of phospho-eIF4E in both the tumor and stromal cells is sufficient to reduce lung metastasis in the PyMT transgenic model of breast cancer (24). To dissect the importance of stromal phospho-eIF4E specifically in PPBC, we investigated whether stromal phospho-eIF4E deficiency is sufficient to block metastasis in a pre-clinical mouse model of this disease. Using the involuting mammary gland as an experimental platform to model PPBC metastasis, 66cl4 murine breast cancer cells were injected into the inguinal mammary glands of wild-type (WT) or eIF4E^S209A/S209A^ (phospho-eIF4E null, henceforth termed eIF4E^S209A^) BALB/c mice one day following weaning-induced involution (Figure 1a). Consistent with previously published data, tumor cells injected into the involuting mammary gland are more metastatic, compared to the same tumor cells injected into virgin mammary glands of age-matched mice (Supplementary Figure 1a). We next harvested the lungs at 33 days post-tumor cell injection for the quantification of metastatic burden. We observed a significant decrease in lung metastases in eIF4E^S209A^ PPBC mice, that is, mice devoid of phosphorylated eIF4E, compared to their WT PPBC counterparts (Figure 1b). The protection of lung metastasis observed in phospho-eIF4E deficient mice was not due to a difference in primary tumor outgrowth, as both tumor initiation, growth and weight at endpoint were similar between WT and eIF4E^S209A^ PPBC mice (Figure 1c, Supplementary Figure 1b and c). Ki67 staining of lung metastases showed no difference in the percentage of Ki67 positive cells in the WT and eIF4E^S209A^ lungs (Figure 1d), indicating similar tumor cell proliferation at the pulmonary metastatic site. Similar results were obtained when E0771 murine breast cancer cells, syngeneic to C57BL/6 mice, were injected into the involuting mammary glands of WT or eIF4E^S209A^ (Figure 1e, 1f, Supplementary Figure 1d, 1e, 1f).

**Figure 1.**
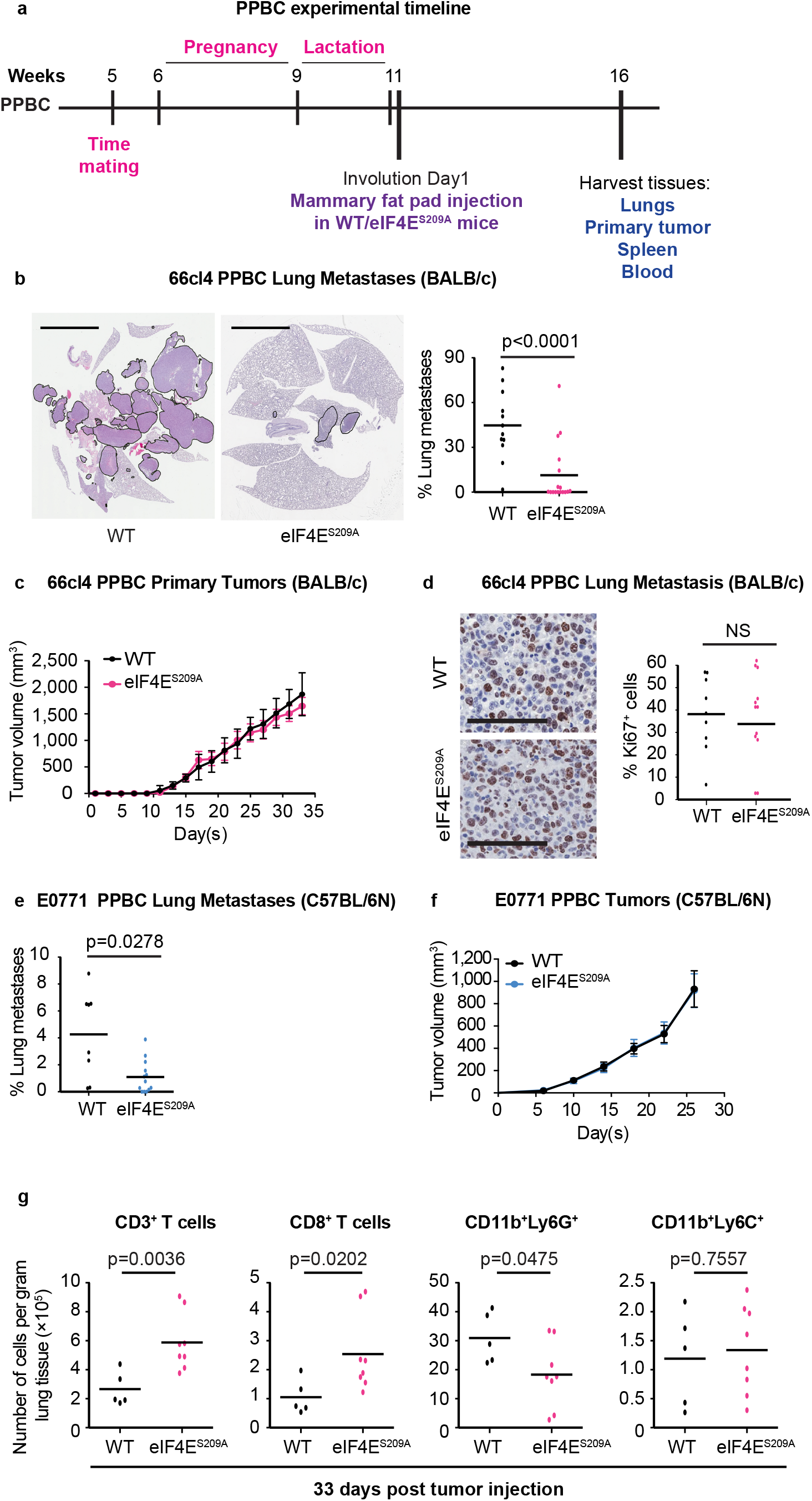
Stromal phospho-eIF4E deficiency (eIF4E^S209A^) protects against lung metastasis in a pre-clinical model of post-partum breast cancer (PPBC) and alters the immune landscape at the lung metastatic site. **a**. Timeline of the PPBC mouse model. **b**. 66cl4 cells were injected into the involuting mammary glands of WT mice or eIF4E^S209A^ phospho-eIF4E deficient PPBC hosts (BALB/c). Less lung metastases are present in eIF4E^S209A^ PPBC mice, compared to WT PPBC mice. Data are presented as average tumor burden in the lung, each dot represents individual mice and the horizontal bar indicates the mean of the cohort. Two-tailed unpaired Student’s t test was used to calculate statistical significance (Scale bar=4mm, WT n=12, eIF4E^S209A^ n=17). **c**. 66cl4 cells injected into in the involuting mammary gland grow at a similar rate in WT and eIF4E^S209A^ PPBC mice. Data are presented as mean tumor volume ±SEM at each time point (WT n=12, eIF4E^S209A^ n=17). **d**. The numbers of Ki67^+^ cells in the lung metastases are similar between the WT and eIF4E^S209A^ BALB/c PPBC mice. Data are presented as % Ki67^+^ cells per lung, each dot represents individual animals. Two-tailed unpaired Student’s t test was used to calculate statistical significance (Scale bar=200μm, WT n=9, eIF4E^S209A^ n=12). **e**. E0771 cells implanted into the involuting mammary glands of phospho-eIF4E deficient PPBC hosts (C57BL/6N) give rise to fewer metastases than in WT PPBC animals. Data are presented as average tumor burden in the lung, each dot represents individual mice and the horizontal bar indicates the mean of the cohort. Two-tailed unpaired Student’s t test was used to calculate statistical significance (WT n=8, eIF4E^S209A^ n=13). **f**. E0771 primary tumors, in the context of the involuting mammary gland, grow at a similar rate in WT and eIF4E^S209A^ C57BL/6N PPBC animals. Data are presented as mean tumor volume ±SEM at each time point (WT n=8, eIF4E^S209A^ n=13). **g**. Immune phenotyping of the lungs of eIF4E^S209A^ PPBC mice injected with 66cl4 tumor cells during mammary gland involution reveals an increased total number of CD3^+^ T cells and cytotoxic CD8^+^ T cells, as well as a decreased number of CD11b^+^ Ly6G cells at 33 days post tumor cell injection. Each dot represents individual mice and the horizontal bar indicates the mean of the cohort. Two-tailed unpaired Student’s t test was used to calculate statistical significance (WT n=5, eIF4E^S209A^ n=8).

Myeloid cells expressing CD11b (marker for myeloid cells of the macrophage lineage) and Gr1 (granulocyte marker present in both Ly6G and Ly6C molecules) are known to increase at pre-metastatic niches and support metastasis (28,33-35). Given the differences in the metastatic burden in the lung, but not primary tumor outgrowth, observed between WT and eIF4E^S209A^ PPBC mice, we focused on whether phospho-eIF4E deficiency at the lung metastatic site altered the infiltration of CD11b^+^, Ly6G^+^, Ly6C^+^ myeloid cell populations. We discovered a significant reduction in granulocytes (CD45^+^CD11b^+^Ly6G^+^Ly6C^lo^) and elevation of CD8^+^ T cells in the lungs of eIF4E^S209A^ PPBC mice (Figure 1g, and Supplementary Figure 2 for gating strategies). However, the pulmonary levels of monocytic cells (CD45^+^CD11b^+^Ly6G^−^Ly6C^hi^) are similar between WT and eIF4E^S209A^ PPBC mice (Figure 1g). In conclusion, phospho-eIF4E deficiency in the host reduces myeloid cells that support metastasis, increases the presence of CD8^+^ T cells, and impairs lung metastasis.

Mammary gland involution is characterized by the elimination of milk-secreting mammary epithelia and the re-population of adipocytes. We next addressed whether the reduced metastatic burden in the lungs of eIF4E^S209A^ PPBC mice was due to a defect in their ability to undergo the physiologic process of mammary gland involution. We quantified the ratio of adipocytes over epithelial cells at lactation day 8, involution days 2, 4, and 6 in WT and eIF4E^S209A^ mice. The adipocyte/epithelium ratio increases in a similar pattern over the course of WT and eIF4E^S209A^ mammary gland involution (Supplementary Figure 1g), and WT and eIF4E^S209A^ show similar gross morphology during involution (Supplementary Figure 1h). Phosphorylation of STAT3 is known to be induced and required for mammary gland involution (36), thus we also examined the levels of phospho-STAT3 in the WT and eIF4E^S209A^ mice, but found no difference in STAT3 phosphorylation (Supplementary Figure 1i). Together, these results suggest that mice devoid of phospho-eIF4E undergo the physiological process of involution similarly to their WT counterparts. Thus, the reduced metastasis observed in phospho-eIF4E null PPBC mice is not the result of overt defects in mammary gland involution.

### Phosphorylated eIF4E regulates IL-33 expression in fibroblasts to support breast cancer cell invasion

Tumor cell invasion and metastasis are regulated to a large degree by molecular signals that can originate within the primary tumor microenvironment (TME). We hypothesized that the reduction in metastatic colonies observed in the lungs of the eIF4E^S209A^ PPBC mice is reflective of a differential expression of such signals in the TME of phospho-eIF4E deficient hosts. Recently, the IL-33/ST2 signaling axis has been implicated as a potent modulator of the TME, regulating the recruitment and activation of immune cells but also tumor cell invasiveness (37-39). We performed IHC on the 66cl4-derived primary tumors that were grown for 2 weeks either in WT or eIF4E^S209A^ PPBC mice, and found that IL-33 levels are lower in the tumors grown in eIF4E^S209A^ PPBC mice, compared to those tumors derived from WT PPBC mice (Figure 2a). Moreover, the expression of phospho-eIF4E correlates with IL-33 expression in WT PPBC tumors (Figure 2b).

**Figure 2.**
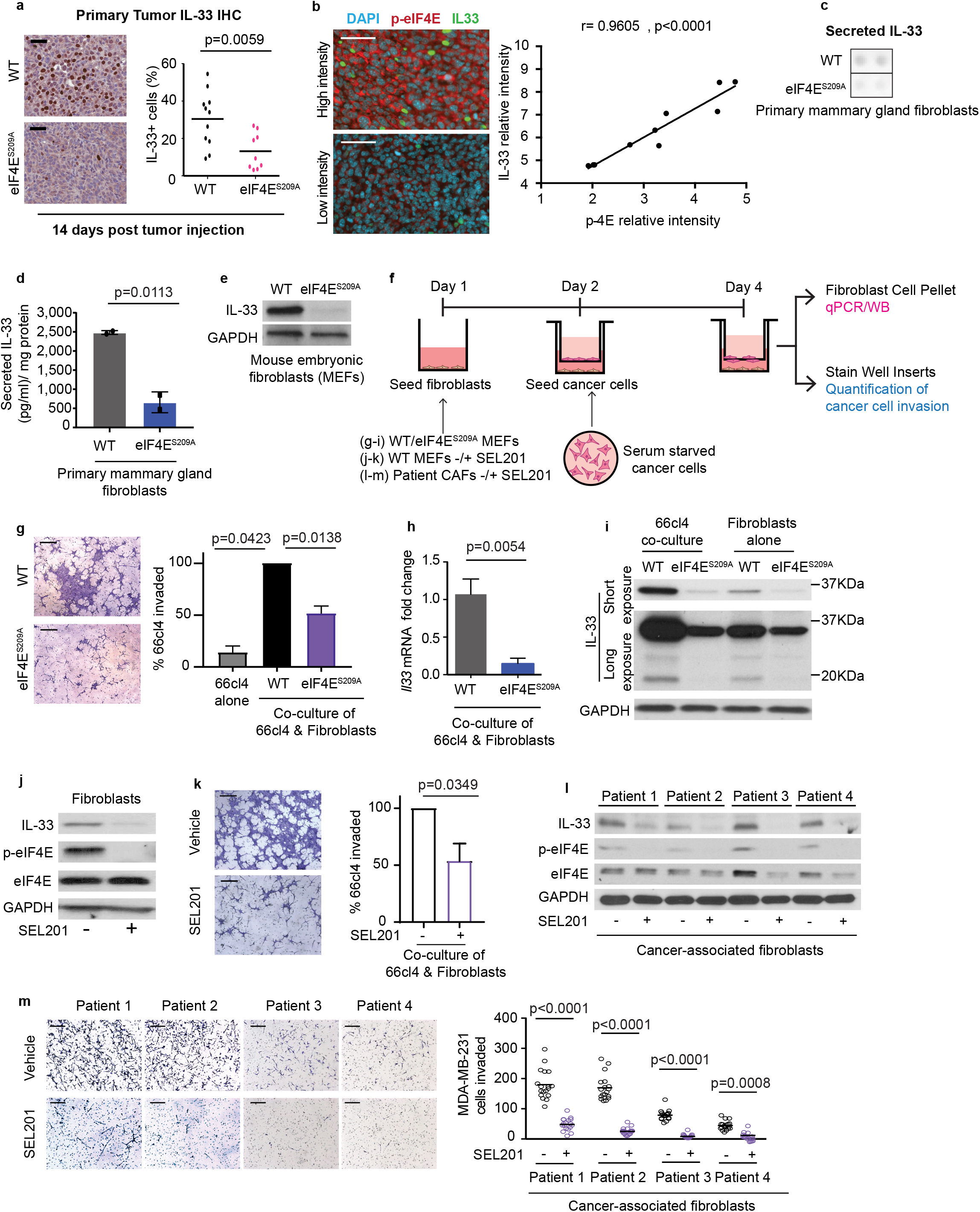
phospho-eIF4E deficient fibroblasts secrete less IL-33 and repress tumor cell invasion in co-culture assays. **a**. 66cl4 PPBC tumors grown in WT BALB/c hosts express more IL-33, than the same tumors grown in phospho-eIF4E (eIF4E^S209A^) deficient hosts (Scale bar=40μm, WT n=11, eIF4E^S209A^ n=9). **b**. The correlation between the expression of phospho-eIF4E (referred to as p-eIF4E in the figures henceforth) and IL-33 in 9 PPBC tumors was assessed by calculating Spearman correlation coefficient. A representative image showing IL-33 and p-eIF4E expression (Scale bar=50μm). **c**. Membrane-based cytokine array profiling of the conditioned media from WT and eIF4E^S200A^ primary mammary gland fibroblasts. Shown is the expression of IL-33 detected in the conditioned media from one representative pool of cells per condition, each pool comprising 3 individual animals. **d**. Cultured primary mammary gland eIF4E^S209A^ fibroblasts secrete less IL-33 than WT mammary fibroblasts. Data represent two separate pools of cells per condition (different from data shown in 3c), each originating from three animals and are presented as mean ±SD. Two-tailed unpaired Student’s t test was used to calculate statistical significance. **e**. eIF4E^S209A^ mouse embryonic fibroblasts express less IL-33. **f**. Schematic overview of the mouse embryonic fibroblast (MEF)-tumor cell co-culture experiments used to generate data in panels **3g-m. g**. 66cl4 cells invade less in the presence of eIF4E^S209A^ fibroblasts, compared to co-culturing with WT fibroblasts. Bars represent the percentage of 66cl4 cells invaded, compared to 66cl4 co-cultured with WT fibroblasts. A summary of three biological replicates is shown. One-way ANOVA was used to calculate statistical significance. **h**. eIF4E^S209A^ fibroblasts express less *Il33* mRNA after co-culture with 66cl4 cells. Data represent three biological replicates normalized to housekeeping gene *Rplp0*) and are presented as mean ±SD. Mann-Whitney test was used to calculate statistical significance. **i**. eIF4E^S209A^ fibroblasts express less IL-33 protein compared to WT fibroblasts, IL-33 expression is induced in WT, but not eIF4E^S209A^, fibroblasts when 66cl4 cancer cells are present. One representative experiment out of three biological replicates is shown. **j**. SEL201 inhibits IL-33 protein expression in WT fibroblasts when co-cultured with 66cl4 cells. One representative experiment out of three biological replicates is shown. **k**. SEL201-treatment of WT fibroblasts suppresses 66cl4 invasion, when co-cultured. Bars represent the percentage of 66cl4 cells invaded compared to 66cl4 co-cultured with WT fibroblasts in absence of SEL201 and a summary of three biological replicates is shown. Two-tailed unpaired Student’s t test was used to calculate statistical significance. **l**. SEL201 treatment decreases IL-33 and p-eIF4E protein levels in patient-derived CAFs. **m**. MDA-MB-231 breast cancer cell invasion is repressed when co-cultured with SEL201-treated patient-derived CAFs. Data represent one experiment per donor. Every dot represents number of cells invaded per field of image (representative images shown). Two-tailed unpaired Student’s t test was used to calculate statistical significance.

Next, we sought to determine the cellular components in the PPBC tumors that produce IL-33. Fibroblasts become activated during mammary gland involution, and they support PPBC invasion and metastasis, in part, via their active secretome (4). We thus sought to determine whether fibroblasts were a major source of IL-33 in our PPBC model. We isolated primary fibroblasts from the mammary glands of WT and eIF4E^S209A^ mice, expanded them *ex vivo* and analyzed their secretome. Mammary fibroblasts derived from eIF4E^S209A^ mice were found to secrete lower levels of IL-33 compared to WT cells (Figure 2c and 2d). We also exploited mouse embryonic fibroblasts (MEFs) derived from WT or eIF4E^S209A^ mice as an additional genetic tool to confirm that the phosphorylation of eIF4E was indeed required for the regulation of IL-33 protein expression (Figure 2e).

IL-33 has been shown to directly impact invasion and metastasis via binding its receptor ST2, which is encoded by interleukin 1 receptor-like 1 (*Il1rl1*) on tumor cells (38,40). Hence, we sought to determine whether fibroblast-derived IL-33 positively supports breast tumor cell invasion. We used a co-culture model system to study interactions between fibroblasts and the 66cl4 and E0771 breast cancer cells used in our *in vivo* PPBC models (Figure 2f). When 66cl4 or E0771 breast cancer cells were co-cultured with WT or eIF4E^S209A^ fibroblasts, both breast cancer cell lines displayed a decreased propensity to invade through a Collagen I matrix in the presence of the eIF4E^S209A^ fibroblasts, as compared to WT fibroblasts (Figure 2g, Supplementary Figure 3a). We also observed a robust increase in the expression of fibroblast-derived IL-33 mRNA and protein when we cultured breast cancer cells in the presence of fibroblasts, however eIF4E^S209A^ fibroblasts still express significantly less IL-33 mRNA and protein, as compared to their WT counterparts (Figure 2h and 2i, Supplementary Figure 3b and 3c).

As we observed that breast cancer cells display an increased propensity to invade toward WT fibroblasts compared to eIF4E^S209A^ fibroblasts, we next examined whether we could pharmacologically inhibit this process using the MNK1/2 inhibitor SEL201. WT fibroblasts were treated with SEL201, and co-cultured with either 66cl4 or E0771 cells. Concomitant with repressed phospho-eIF4E expression in fibroblasts, SEL201 treatment decreased IL-33 protein levels in WT fibroblasts (Figure 2j, Supplementary 3d). Moreover, the invasion of 66cl4 and E0771 cells was less robust when co-cultured with SEL201-treated fibroblasts (Figure 2k, Supplementary Figure 3e).

Finally, we addressed the clinical relevance of our findings by co-culturing patient-derived CAFs with MDA-MB-231 human breast cancer cells. We obtained primary CAFs that were isolated from the freshly resected human breast tumors of four patients. Primary CAFs were treated with vehicle or SEL201, and subsequently co-cultured with MDA-MB-231. Similar to our findings in the murine fibroblasts, SEL201 repressed phospho-eIF4E and IL-33 expression in patient-derived CAFs, and MDA-MB-231 invaded less robustly in the presence of SEL201-treated CAFs (Figure 2l and 2m). Collectively, our data show that IL-33 expression in fibroblasts is regulated by the MNK1/2-eIF4E axis.

### IL-33 activates the MNK1/2-eIF4E pathway downstream of activated ST2 to support an immune suppressed TME

Having shown the important role of fibroblast-derived IL-33 in supporting breast cancer cell invasion, as well as the reduction of IL-33 in the TME of eIF4E^S209A^ PPBC tumors, we next investigated whether fibroblast-derived IL-33 acts via the IL-33 receptor, ST2, expressed on 66cl4 cells, to promote breast cancer invasion. By ablating the expression of *Il1rl1* using siRNA in 66cl4 cells, we observed an impaired ability of the ST2-deficient tumor cells to invade in the presence of WT fibroblasts (Figure 3a, Supplementary Figure 4a). Such data indicate that fibroblast-derived IL-33 signals in a paracrine fashion to ST2-expressing breast cancer cells to augment tumor cell invasion. Therefore, we chose to further dissect how IL-33 signals downstream of ST2 in breast tumor cells, focusing on the p38 and ERK1/2 MAPK signaling proteins, which lie immediately upstream of MNK1/2 activation(41). Stimulation of 66cl4 cells with recombinant murine IL-33 (rIL-33) resulted in increased phosphorylation of both p38 MAPK and eIF4E, but not phosphorylation of ERK1/2 (Figure 3b, Supplementary 4b), and promoted 66cl4 (Figure 3c) and E0771 invasion (Supplementary Figure 4c). Additionally, we hypothesized that IL-33 might stimulate the expression of pro-inflammatory and pro-tumorigenic cytokines/chemokines in tumor cells, which may further remodel the TME to favor invasion. rIL-33 stimulated the mRNA expression of *Cxcl1, Ccl17, Csf2* (which encodes GM-CSF) and *Il6*, without significantly affecting *Il4* and *Cxcl2* levels (Figure 3d). Finally, a main immune cell type whose expansion is reliant on IL-33/ST2 signaling, and which have been shown to play an important role in tumor immunity, are ILC2 cells (10). Using *ex vivo* expanded ILC2s from the bone marrow of BALB/c mice, we examined their cellular viability as well as their ability to secrete IL-5 and IL-13 in response to the co-stimulation with rIL-7 plus rIL-33 in the presence and absence of SEL201 (42). SEL201 treatment of ILC2s reduced their secretion of IL-5 and IL-13, in response to combined rIL-7 and rIL-33 (Figure 3e), with minimal effects on their cell viability (Figure 3f). Together, these results suggest that the phosphorylation of eIF4E is necessary for ILC2-derived IL-5 and IL-13, which are important for the recruitment of CD11b^+^Gr1^+^ cells (9-12,43-46). In toto, given the reported functions of CXCL-1, CCL-17, GM-CSF, IL-6, IL-5 and IL-13 in tumor immune evasion (44,45,47-55), our data provide evidence that IL-33 may serve to create an immunosuppressive PPBC TME to facilitate metastasis in a MNK1/2-phospho-eIF4E-dependent way.

### Characterization of the human PPBC TME

The expression of phospho-eIF4E positively correlates with IL-33 protein level in 66cl4 tumors grown in WT PPBC mice (Figure 2b). To interrogate whether these observations are clinically relevant, we used immunofluorescence (IF) staining to evaluate the expression of phospho-eIF4E and IL-33 in a cohort of PPBC patient samples. Consistent with our PPBC murine data, phospho-eIF4E expression correlates with IL-33 levels in human PPBC tumors (Figure 4a). To verify the broader implications of our observations, we examined TCGA data using UCSC Xena (http://xenabrowser.net/) for the relationship between MNK1 and IL-33 expression. We found that *MKNK1* and *IL33* mRNA levels significantly correlate with one another (Figure 4b).

**Figure 3.**
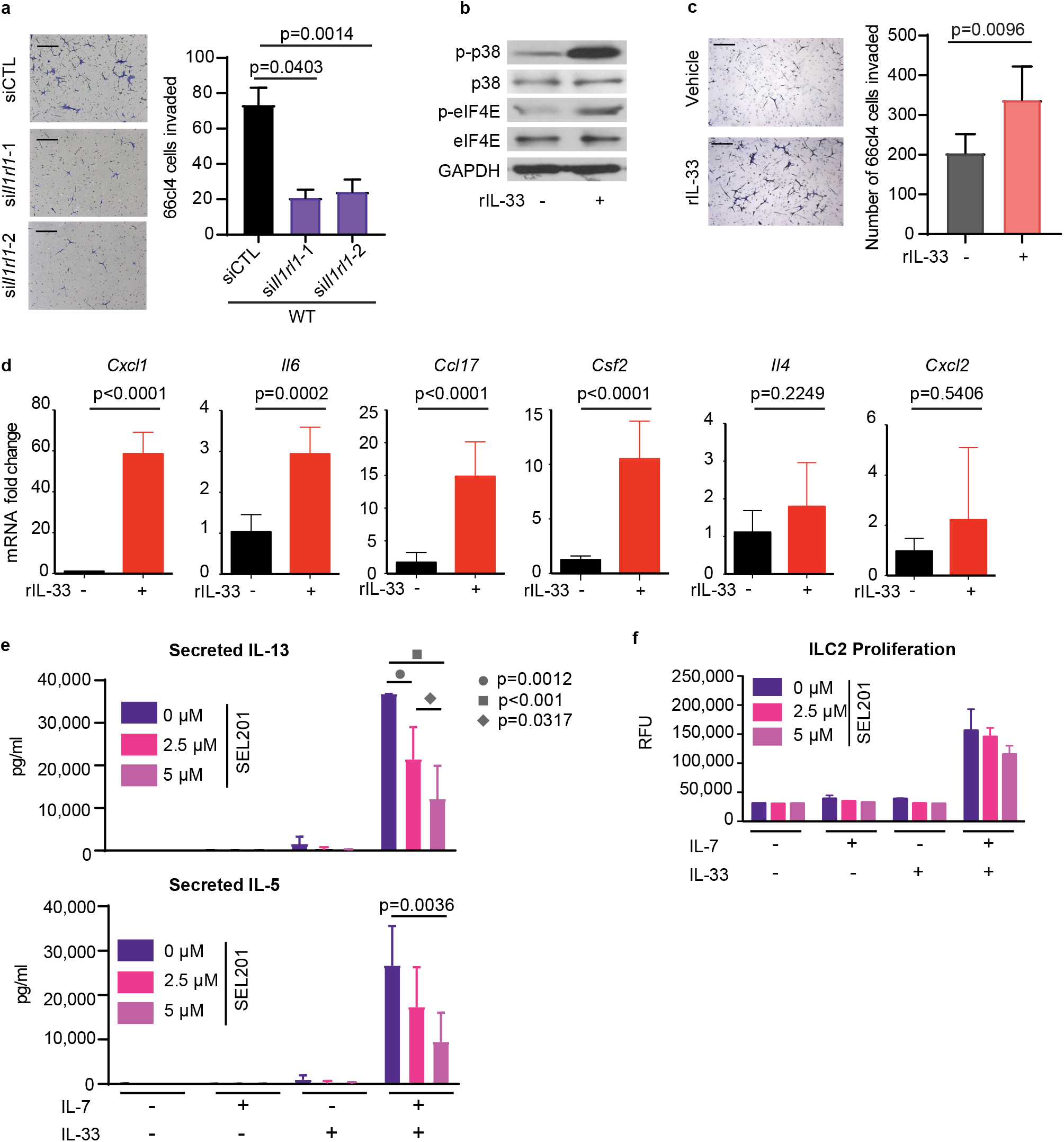
Recombinant IL-33 induces tumor cell invasion and upregulates the expression of immune suppressive chemokines and cytokines. **a**. Knockdown of *Il1rl1* (encodes ST2) in 66cl4 cells diminishes their invasion toward fibroblasts. Bars represent the number of cells invaded per field of image (representative images shown) and a summary of three biological replicates is shown. One-way ANOVA was used to calculate statistical significance. **b**. 50ng/ml rIL-33 treatment for 6hrs induces the phosphorylation of p38 and eIF4E in 66cl4 cells. One representative experiment out of three is shown. **c**. rIL-33 enhances 66cl4 cell invasion. Bars represent the number of cells invaded per field of image and a summary of three biological replicates is shown. The horizontal bar indicates the mean. Mann-Whitney test was used to calculate statistical significance. **d**. rIL-33 induces the expression of *Cxcl1, Ccl17, Csf2*, and *Il6* mRNA in 66cl4 cells. rIL-33 does not induce the mRNA expression of *Il4* and *Cxcl2* in 66cl4 cells. Data represent three biological replicates and are presented as mean ±SD. Mann-Whitney test was used to calculate statistical significance. **e**. SEL201 treatment reduces IL-13 and IL-5 secretion of WT ILC2s. Two-way ANOVA was used to calculate significance **f**. SEL201 does not impact the proliferation of WT ILC2s. Panels (**e-f**), ILC2 cells were isolated from the bone marrow of BALB/c mice and ex vivo expanded. Data in **(e-f)** represent two biological replicates, each comprising 3 technical replicates and are presented as mean ± SD.

**Figure 4.**
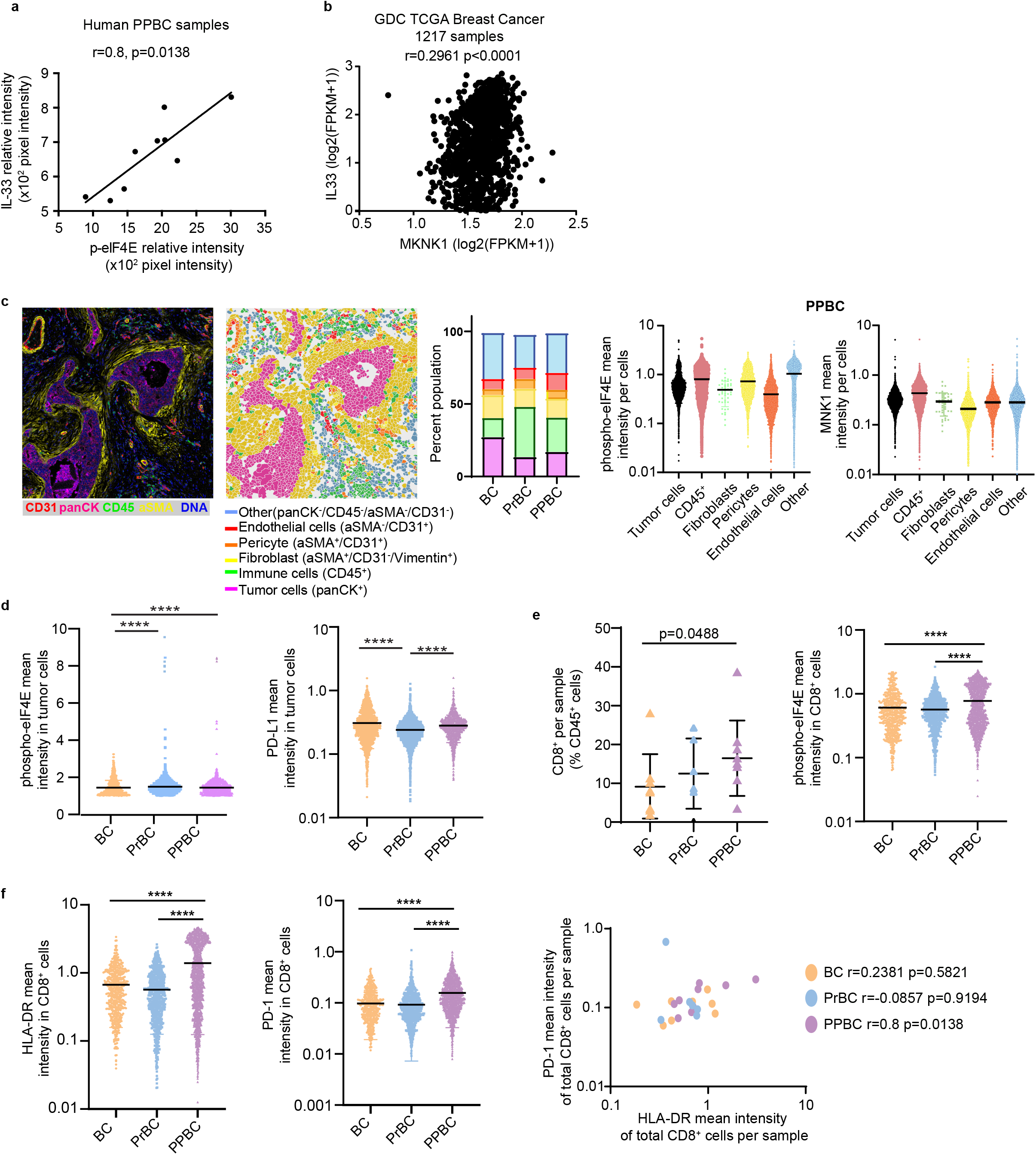
Imaging of the human PPBC TME. **a**. Positive correlation between IL-33 protein expression in human PPBC samples (n=9) and p-eIF4E levels. Spearman correlation was used to calculate statistical significance. **b**. Spearman correlation analysis of the mRNA expression of *MKNK1* and *IL33* using the TCGA breast cancer database (1217 samples). **c**. Imaging mass cytometry (one image shown, left) revealed the proportion of different cell types, and their level of phospho-eIF4E expression, in 8 human nulliparous breast tumors (BC), 6 pregnancy associated breast cancers (PrBC), and 9 postpartum breast cancers (PPBC). Each dot represents one cell. One-way ANOVA was used to calculate statistical significance. **d**. Phospho-eIF4E and PD-L1 expression are detected in human tumor cells. Each dot represents one cell. One-way ANOVA was used to calculate statistical significance. **e**. PPBC samples have increased CD8^+^ T cell infiltration, compared to BC and PrBC samples. Each dot represents the proportion of cell per core. One-way ANOVA was used to calculate statistical significance. **f**. CD8^+^ T cells in PPBC have increased surface expression of HLA-DR and PD-1, compared to BC and PrBC. Each dot represents one cell. One-way ANOVA was used to calculate statistical significance. Expression of CD8^+^HLA-DR^+^ T cells and CD8^+^PD-1^+^ T cells correlate in each individual PPBC core. Each dot represents the proportion of cell per core. Spearman correlation was used to calculate statistical significance.

The limited success of immune targeted therapy in breast cancer, relative to other malignancies, has been attributed in part to the heterogeneity of the breast TME. The TME of human PPBC has not been well defined, and we used CyTOF imaging mass cytometry (IMC) to simultaneously quantify the expression of 26 proteins within the TME of nulliparous breast cancer (BC), breast cancer diagnosed during pregnancy (PrBC), and postpartum breast cancer (PPBC) (Supplementary Table 4 and Supplementary Figure 5a). We report that PrBC and PPBC differ from BC primarily in the relative proportion of tumor cells and immune cells, as well as their level of activation of the MNK1/2-eIF4E axis. Phospho-eIF4E, eIF4E, and MNK1 were detectable in tumor cells, immune cells, fibroblasts, pericytes, and endothelial cells (Figure 4c and Supplementary Figure 5b, c and d). In particular, the tumor cell population in PPBC and PrBC showed a significantly increased level of phospho-eIF4E expression, compared to the tumor cells represented in BC. Furthermore, PD-L1 was expressed in tumor cells regardless of the cancer subtype, with PD-L1 expression being significantly increased in PPBC tumor cells compared to PrBC tumor cells (Figure 4d). Moreover, when we examined tumor cells with the highest expression of phospho-eIF4E (99^th^ percentile), the expression of PD-L1 was most abundant in PPBC, compared to BC and PrBC (Supplementary Figure 5e). CD8 is an important tumor immune biomarker, so we next investigated the proportion of CD8^+^ T cells in the three patient cohorts. We observed a significant increase in CD8^+^ T cells in the PPBC samples, and the phosphorylation of eIF4E is significantly increased in the CD8^+^ T cells present in PPBC, compared to BC or PrBC samples (Figure 4e). Further characterization of the phenotype of the CD8^+^ T cells showed that the co-expression of HLA-DR and PD-1 was significantly higher in PPBC samples, compared to CD8^+^ T cells present in PrBC or BC (Figure 4f). As HLA-DR is known to be expressed on activated T cells, and PD-1 is an exhaustion marker, our data suggest that the CD8^+^ T cells present in PPBC express an activated dysfunctional phenotype (56,57).

### Dual MNK1/2 and PD-1 blockade inhibits PPBC lung metastasis

The efficacy of immune checkpoint inhibitors in metastatic breast cancer, including PPBC, would likely be improved by overcoming anti-tumor immune escape. Given the diverse roles of phospho-eIF4E in contributing to PPBC immune evasion, we hypothesized that SEL201 might sensitize PPBC mice to the anti-metastatic effects of PD-1 blockade. To test this hypothesis, WT PPBC mice were administered vehicle, SEL201, anti-PD-1 antibody, or the combination of SEL201 plus anti-PD1 antibody (Supplementary Figure 5f). Remarkably, SEL201 plus anti-PD-1 blockade decreased PPBC lung metastasis, while SEL201 or anti-PD-1 alone did not show any significant anti-metastatic effects (Figure 5a). SEL201 treatment resulted in a significant decrease in phospho-eIF4E expression, suggesting efficient target engagement by the MNK1/2 inhibitor, and a robust increase in CD8^+^ cells in the lung metastases (Figure 5b, Supplementary Figure 5g). The reduction in metastatic burden in the mice treated with the combination of SEL201 and anti-PD-1 was not due to a difference in the effect of the combination therapy on primary tumor outgrowth, which remained unchanged (Figure 5c, Supplementary Figure 5h). Administration of SEL201 or anti-PD-1 did not have any overt systemic toxicity (Supplementary Figure 4i), consistent with our previous work (23,26). Targeting the MNK1/2-eIF4E axis might have therapeutic benefit for augmenting the therapeutic efficacy of immunotherapy in women diagnosed with PPBC.

**Figure 5.**
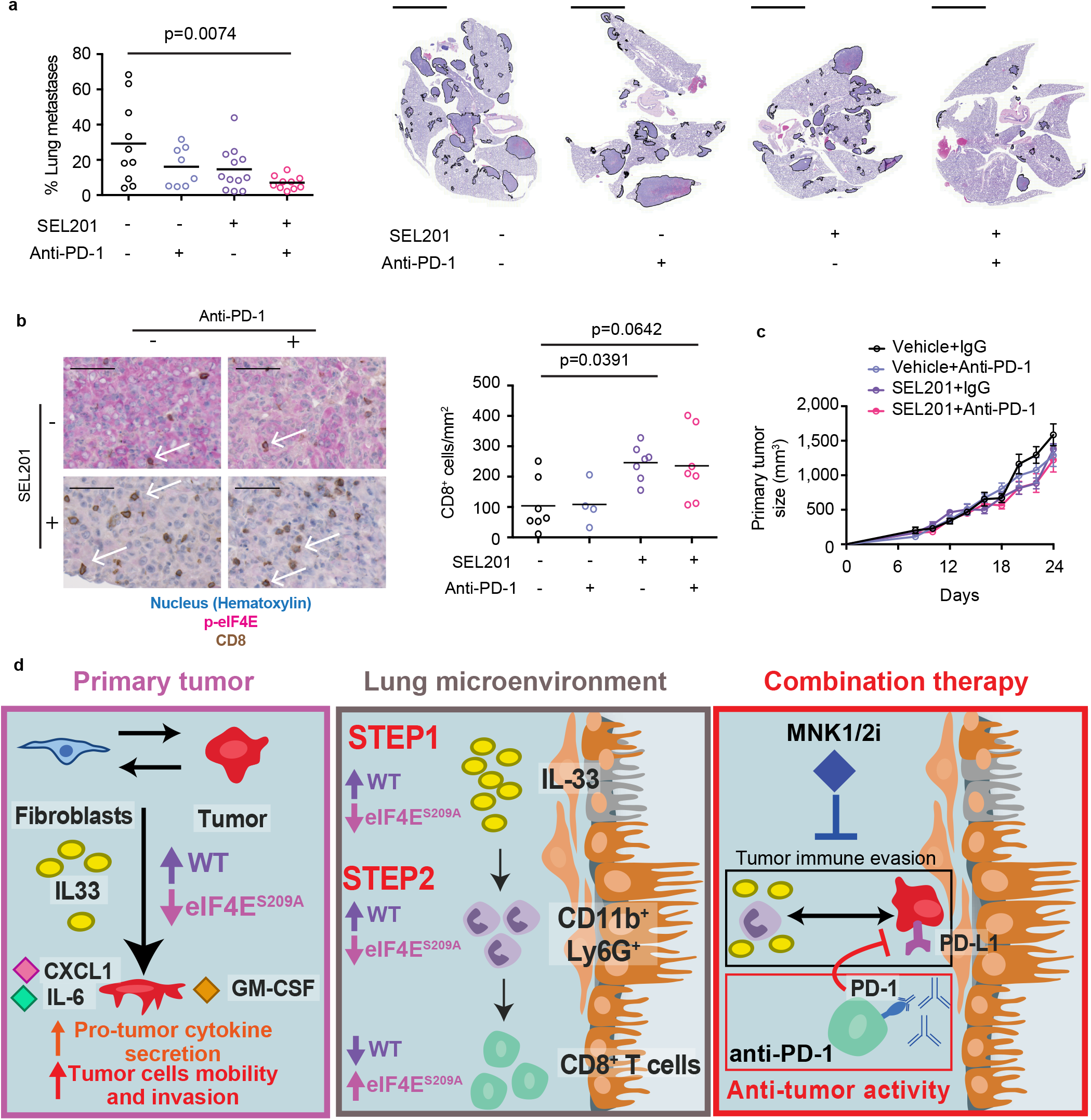
Sensitization of murine PPBC to PD-1 blockade using a MNK1/2 inhibitor. **a**. Combination of SEL201 plus anti-PD1 significantly decreased 66cl4 lung metastatic burden in the PPBC model. Data are presented as average tumor burden in the lung (representative images of the lung metastases are shown), each dot represents individual animals and the horizontal bar indicates the mean of the cohort. One-way ANOVA was used to calculate statistical significance. Scale bar=4mm. Vehicle group n=10, anti-PD-1 treated group n=8, SEL201-treated group n=12, SEL201+anti-PD1 group n=10. **b**. SEL201 treatment efficiently repressed phospho-eIF4E expression and significantly increased the number of CD8^+^ T cells in the lung metastases (shown by IHC). White arrows on the representative images indicate CD8^+^ T cells. Scale bar=50μm. Each dot in the graphs represents one animal and the horizontal bar indicates the mean of the cohort. One-way ANOVA was used to calculate statistical significance. Vehicle group n=7, anti-PD-1 treated group n=5, SEL201-treated group n=7, SEL201+anti-PD1 group n=7. **c**. SEL201 administered alone or in combination with anti-PD-1 blockade does not significantly alter primary tumor outgrowth. Data are presented as mean tumor volume ±SEM at each time point. Vehicle group n=10, anti-PD-1 treated group n=8, SEL201-treated group n=12, SEL201+anti-PD-1 group n= 10. **d**. Summary of the pleiotropic effects mediated by the MNK1/2-eIF4E axis to support immune suppression and PPBC metastasis.

## Discussion

Metastasis associated with PPBC and mortality due to lack of effective treatment strategies necessitate a fuller understanding of this disease (58,59). Recent breakthroughs in immune checkpoint blockade therapies have stimulated research to better understand the TME of breast cancer, aiming to discover possible approaches to sensitize metastatic breast cancer to immunotherapies. Here, we demonstrated the central role of stromal phospho-eIF4E in promoting pro-tumorigenic immunity in a model of metastatic PPBC (Figure 5d, graphic summary). Collectively, our work highlights the important role of the IL-33-MNK1/2-eIF4E axis in PPBC invasion and metastasis by impacting multiple cellular compartments in the TME. IL-33 might hold potential as a therapeutically targetable cytokine in PPBC, and perhaps more broadly in breast cancer.

The regulation of IL-33 by the MNK1/2-eIF4E axis is potentially relevant for immune cell function. IL-33 has important effects on numerous immune cell types such as eosinophils, mast cells and ILC2s (5,6). IL-33 is essential for the polarization of ILC2s together with IL-7, IL-25 and TSLP (60). Our understanding of the roles ILC2 plays in the context of tumor biology is still rudimentary, although recent studies have described their tumor-promoting and anti-tumor roles in several cancer types, including breast cancer (11,12,61,62). In our study, we showed that the pharmacologic inhibition of MNK1/2 blocked the IL-33 induced expression of *ex vivo* expanded ILC2-derived cytokines IL-5 and IL-13. We have yet to test whether hosts devoid of phospho-eIF4E, which presented with less granulocytic CD45^+^CD11b^+^Ly6G^+^Ly6C^lo^ cells in the lungs, is due to suppression of the IL-33-ILC2-MDSC axis (12).

An additional function of IL-33 is to reinforce pro-tumorigenic inflammation by inducing IL-6 (40). Our study has not only verified this finding in breast cancer, but also expanded the repertoire of IL-33-induced cytokines and chemokines (i.e. *Cxcl1, Ccl17, Il6* and *Csf2*) produced by tumor cells. The significance of these four factors in tumor immune evasion has been supported by multiple previous reports. For example, over-expression of CXCL-1 and its receptor CXCR-2, as well as elevated circulating IL-6 levels, are all correlated to breast cancer metastasis and poor survival rate (63,64), and CXCL-1, IL-6 and GM-CSF are all potent mediators for the recruitment and expansion of MDSCs and M2-like macrophages (48-51,53-55). CCL-17, an important ligand for CCR-4, has also been demonstrated to elicit Th2 and T_reg_-mediated cancer immune evasion (47,52). Taken together, we show that IL-33 acts directly on breast tumor cells to induce the expression of selected immunosuppressive chemokines and cytokines.

In addition to its impact on immune cells, IL-33 has also been reported as a pro-tumorigenic and pro-invasive cytokine (37-39). Elevation of IL-33 was observed in the serum of breast cancer patients (65,66). Moreover, the levels of matrix metallopeptidase 11 (*MMP11*), a pro-invasive enzyme responsible for tissue remodeling, are directly correlated with *IL33* levels in patients with breast cancer (66), supporting a possible pro-invasive function of IL-33. In this context, we have demonstrated that phospho-eIF4E-mediated IL-33 production in fibroblasts is further induced when fibroblasts are co-cultured with breast tumor cells. Disrupting IL-33/ST2 signaling diminished the ability of cancer cells to invade. Thus, our data highlight the crosstalk between cancer cells and fibroblasts, whereby cancer cells educate fibroblasts to secrete more IL-33, thus allowing breast cancer cells to gain more invasive properties and implicate the MNK1/2-phospho-eIF4E axis as the driver of this crosstalk.

Finally, immune checkpoint blockades designed to release the brakes on exhausted cytotoxic T cells have largely improved the patient prognosis in several cancers (67), but are less effective to date in breast cancer. It is proposed that many breast cancers display failed or suboptimal T cell priming. Specifically in PPBC, increase in PD-1 expression on T cells and efficacy of anti-PD-1 treatment in reversing involution-associated tumor growth was recently reported (68). In line with this finding, we observe increased expression of PD-1 on the tumor-infiltrating CD8^+^ cells in PPBC patients. We propose that in the TME of PPBC tumors, these cells present a dysfunctional immunosuppressive phenotype. In this context, it is interesting that these CD8^+^ cells also express elevated levels of phospho-eIF4E. While the role of phospho-eIF4E in regulating functionality of specifically CD8^+^ has not yet been explored by the scientific community and is beyond the scope of this present study, emerging data suggest that the activity of MNK1/2-eIF4E axis in immune cells affects their function. For example, a recent paper reported that the immunosuppressive phenotype of bone-marrow derived macrophages is governed by the MNK1/2-eIF4E axis and can be reversed by MNK2 inhibition, indirectly leading to increased CD8^+^ cell activation (Bartish, Tong et al, PNAS 2020 *in press*). Furthermore, a pre-clinical study of the MNK1/2 inhibitor Tomivosertib (eFT508) in liver cancer has shown that this inhibitor enhances the activity of checkpoint inhibitors in a T-cell dependent manner, leading to an anti-tumor immune response (69). In summary, we propose that MNK1/2 inhibitors may convert “cold” breast tumors to “hot” tumors, thus offering the opportunity for immune checkpoint blockade to become more effective in highly metastatic cancers such as PPBC. This study, as well as others showing effects of MNK1/2 inhibition on cells of the TME (28), provide strong preclinical support to ongoing clinical testing of MNK1/2 inhibitors in breast cancer. Indeed, we are participating in a Stand Up to Cancer trial to test this question (ClinicalTrials.gov NCT04261218).

## Authors’ Contributions

*Conception and design*: Qianyu Guo, Wilson H. Miller Jr., Claudia U. Duerr, Jörg H. Fritz, Sonia V. del Rincón

*Development of methodology*: Qianyu Guo, Margarita Bartish, Fan Huang, Sai Sakktee Krisna, Audrey Emond, Claudia U. Duerr, Christophe Goncalves, Sonia V. del Rincón

*Acquisition of data*: Qianyu Guo, Margarita Bartish, Fan Huang, Sai Sakktee Krisna, Samuel E. J. Preston, Vivian Zihui Li, Hanne Lefrère, Audrey Emond, Claudia U. Duerr, Barbara C. Mindt, Yirui Gui, Christophe Goncalves, Dany Plourde, Jie Su, Shannon Hewgill, William Yang, Elie Khoury, Yao Zhan, Valeria Narykina

*Analysis and interpretation of data*: Qianyu Guo, Margarita Bartish, Christophe Goncalves, Sai Sakktee Krisna, Audrey Emond, Samuel E. J. Preston, Aurélie Cleret-Buhot, Wilson H. Miller Jr.

*Writing, review, and/or revision of the manuscript*: Qianyu Guo, Margarita Bartish, Sonia V. del Rincón, Wilson H. Miller Jr.

*Administrative, technical, or material support:* Mark Basik, Paméla Thébault, Tiziana Cotechini, Charles Hindmarch, Liesbeth Lenaerts, Yuhong Wei, Frédéric Amant, Daniela F. Quail, Réjean Lapointe, Jörg H. Fritz, Giuseppe Floris

*Study supervision:* Qianyu Guo, Wilson H. Miller Jr., Sonia V. del Rincón

## Acknowledgements

We thank Christian Young (LDI Flow Cytometry facility), Dr. Naciba Benlimame, and Lilian Canetti for experimental advice and technical support. We thank Dr. Pepper Schedin (Oregon Health & Science University) and Dr. Qiuchen Guo (Harvard Medical School) for helpful discussion and experimental design.

