## Supplemental Table for "The pleiotropic effects of the MNK1/2-eIF4E axis support immune suppression and metastasis in a model of postpartum breast cancer"

**Supplementary Table 1: Antibodies used for flow cytometry**

| Target | Fluorochrome | Clone | Company |
|---|---|---|---|
| **CD11b** | e450 | M1/70 | eBioscience |
| **CD3e** | BV650 | 145-2C11 | BD Bioscience |
| **CD11c** | BV786 | HL3 | BD |
| **Ly6G (Gr1)** | AF488 | RB6-8C5 | eBioscience |
| **CD8a** | PerCP Cy5.5 | 53-6.7 | BD |
| **F4/80** | PE | BM8 | eBioscience |
| **Ly6C** | APC | AL-21 | BD |
| **CD45** | APC-Cy7 | 30-F11 | BD |
| **Fixable viability dye** | Aqua | N/A | Invitrogen |
| **Gr1** | e450 | RB6-8C5 | eBioscience |
| **PDL-1** | APC | MIH5 | BD |

### Supplementary Table 2: Antibodies used for IF, IHC and WB

| Target protein | Antibody information | Usage |
|---|---|---|
| **α-SMA** | #18147 Abcam | IF |
| **eIF4E** | #610269 BD Transduction | WB |
| **eIF4E** | #2067 Cell Signaling | IHC, IF |
| **phospho-eIF4E** | #9741 Cell Signaling | WB |
| **phospho-eIF4E** | #ab76256 Abcam | IHC, IF |
| **MNK1** | #2195 Cell Signaling | WB, IHC, IF |
| **phospho-p38** | #4511 Cell Signaling | WB |
| **p38** | #sc-535-G Santa Cruz | WB |
| **phospho-ERK1/2** | ##4370 Cell Signaling | WB |
| **ERK1** | #sc-93 Santa Cruz | WB |
| **ERK2** | #sc-154 Santa Cruz | WB |
| **IL-33 (mouse)** | #AF3626 R&D System | WB/IHC/IF |
| **IL-33 (human)** | #AF3625 R&D System | WB/IHC/IF |
| **CD8a** | #14-0808 thermofischer | IHC |
| **Vimentin** | #550513 BD Pharmingen | WB |
| **Ki67** | #15580 Abcam | IHC |
| **GAPDH** | #2118 Cell Signaling | WB |

**Supplementary Table 3: Details of qPCR Primers (Mouse)**

| Target gene | Primer Sequence |
|---|---|
| ***Il33*** | Fwd: 5’-ATG GGA AGA AGC TGA TGG TG-3’ |
| | Rev: 5’-CCG AGG ACT TTT TGT GAA GG-3’ |
| ***Cxcl1*** | Fwd: 5’-CAC CTC AAG AAC ATC CAG AGC-3’ |
| | Rev: 5’-CTT GAG TGT GGC TAT GAC TTC G-3’ |
| ***Cxcl2*** | Fwd: 5’-CATCCAGAGCTTGAGTGTGACG-3’ |
| | Rev: 5’-GGCTTCAGGGTCAAGGCAAACT-3’ |
| ***Il4*** | Fwd: 5’-ATCATCGGCATTTTGAACGAGGTC-3’ |
| | Rev: 5’-ACCTTGGAAGCCCTACAGACGA-3’ |
| ***Il6*** | Fwd: 5’-CAT GTT CTC TGG GAA ATC GTG-3’ |
| | Rev: TTC TGC AAG TGC ATC ATC G-3’ |
| ***Ccl17*** | Fwd: 5’-GGA AGT TGG TGA GCT GGT ATA A-3’ |
| | Rev: 5’-GAT GGC CTT CTT CAC ATG TTT G-3’ |
| ***Csf2*** | Fwd: 5’-GAA GAT ATT CGA GCA GGG TCT AC-3’ |
| | Rev: 5’-CTT GTG TTT CAC AGT CCG TTT C-3’ |
| ***Rplp0*** | Fwd: 5’-TCA TCC AGC AGG TGT TTG ACA-3’ |
| | Rev: 5’-GGC ACC GAG GCA ACA GTT-3’ |

**Supplementary table 4: Clinical details of the patient cohort analyzed by imaging mass cytometry**

| Study Group | Age at Diagnosis | ER | PR | HER2 | Molecular Subtype | Stage | Grade | Ki67 | Overall Survival (0=censored, 1=death) | Time to OS (Years) | Time to metastasis (Years) |
|---|---|---|---|---|---|---|---|---|---|---|---|
| **non-PrBC** | 38 | 1 | 1 | 0 | Luminal B (HER2-) | Stage IIIC | Grade III | 15% | 1 | 3,3 | 1,3 |
| **non-PrBC** | 37 | 1 | 1 | 0 | Luminal B (HER2-) | Stage IA | Grade III | x | 0 | 18,6 | 18,6 |
| **non-PrBC** | 26 | 0 | 0 | 0 | Triple Negative | Stage IIIA | Grade III | x | 1 | 9 | 1,1 |
| **non-PrBC** | 26 | 1 | 1 | 0 | Luminal B (HER2-) | Stage IIA | Grade III | x | 0 | 11,8 | 11,8 |
| **non-PrBC** | 40 | 1 | 1 | 0 | Luminal B (HER2-) | Stage IIA | Grade III | x | 0 | 11,1 | 11,1 |
| **non-PrBC** | 38 | 1 | 1 | 0 | Luminal B (HER2-) | Stage IIIA | Grade III | x | 1 | 9,7 | 7,6 |
| **non-PrBC** | 39 | 1 | 1 | 0 | Luminal B (HER2-) | Stage IIA | Grade III | x | 0 | 7,4 | 7,4 |
| **non-PrBC** | 35 | 0 | 0 | 0 | Triple Negative | Stage IIA | Grade III | x | 0 | 23,8 | 23,8 |
| **non-PrBC** | 35 | 0 | 0 | 1 | HER2+ (non-luminal) | Stage IIIA | Grade III | x | 1 | 4,4 | 2,7 |
| **PPBC** | 31 | 1 | 1 | 0 | Luminal B (HER2-) | Stage IIB | Grade III | 70% | 0 | 19,7 | 19,7 |
| **PPBC** | 31 | 0 | 0 | 0 | Triple Negative | Stage IIIB | Grade III | x | 0 | 17,9 | 17,9 |
| **PPBC** | 35 | 1 | 1 | 0 | Luminal B (HER2-) | Stage IIIA | Grade III | x | 1 | 4,1 | 3,2 |
| **PPBC** | 31 | 0 | 0 | 0 | Triple Negative | Stage IA | Grade III | x | 0 | 13,4 | 13,4 |
| **PPBC** | 41 | 1 | 1 | 0 | Luminal B (HER2-) | Stage IB | Grade III | x | 0 | 9,2 | 9,2 |
| **PPBC** | 27 | 1 | 1 | 0 | Luminal B (HER2-) | Stage IA | Grade III | x | 0 | 8,9 | 8,9 |
| **PPBC** | 34 | 1 | 1 | 0 | Luminal B (HER2-) | Stage IIIC | Grade III | x | 0 | 1,6 | 1,6 |
| **PPBC** | 38 | 0 | 0 | 1 | HER2+ (non-luminal) | Stage IIIB | Grade III | 40% | 0 | 4,7 | 4,7 |
| **PPBC** | 38 | 1 | 1 | 0 | Luminal A | Stage IIIA | Grade II | x | 1 | 15,5 | 9,9 |
| **PrBC** | 30 | 0 | 0 | 0 | Triple Negative | Stage IIB | Grade III | x | 0 | 19,0 | 19,0 |
| **PrBC** | 32 | 0 | 0 | 0 | Triple Negative | Stage IA | Grade III | x | 0 | 15,9 | 15,9 |
| **PrBC** | 38 | 0 | 0 | 0 | Triple Negative | Stage IIA | Grade III | x | 0 | 12,2 | 12,2 |
| **PrBC** | 40 | 1 | 0 | 1 | Luminal B (HER2+) | Stage IIA | Grade III | x | 0 | 11,1 | 11,1 |
| **PrBC** | 36 | 0 | 0 | 0 | Triple Negative | Stage IIIA | Grade III | x | 0 | 9,1 | 9,1 |
| **PrBC** | 28 | 0 | 0 | 0 | Triple Negative | Stage IIB | Grade III | x | 0 | 8,0 | 8,0 |
| **PrBC** | 33 | 1 | 1 | 0 | Luminal B (HER2-) | Stage IIB | Grade III | x | 0 | 3,4 | 3,4 |

### Supplementary Table 5: Antibodies used for imaging mass cytometry

| Target | Fluorochrome | Company |
|---|---|---|
| **CD56** | 142Nd | SCIMAP core |
| **Vimentin** | 143Nd | QCPU core |
| **CD14** | 144Nd | SCIMAP core |
| **CD66b** | 145Nd | SCIMAP core |
| **CD16** | 146Nd | SCIMAP core |
| **CD31** | 147Sm | SCIMAP core |
| **Pan-CK** | 148Nd | QCPU core |
| **CD11b** | 149Sm | SCIMAP core |
| **PD-L1** | 150Nd | SCIMAP core |
| **MNK1** | 151Eu | 2195 Cell Signaling |
| **CD45** | 152Sm | SCIMAP core |
| **CD11c** | 154Sm | SCIMAP core |
| **FocP3** | 155Gd | SCIMAP core |
| **CD4** | 156Gd | SCIMAP core |
| **eIF4E** | 158Gd | 2067 Cell Signaling |
| **CD68** | 159Tb | SCIMAP core |
| **CD20** | 161Dy | SCIMAP core |
| **CD8a** | 162Dy | SCIMAP core |
| **Phospho-eIF4E** | 164Dy | ab76256 Abcam |
| **PD1** | 165Ho | SCIMAP core |
| **CD40** | 168Er | SCIMAP core |
| **CD3** | 170Er | SCIMAP core |
| **HLA-DR** | 174Yb | SCIMAP core |
| **a-SMA** | 175Lu | SCIMAP core |
| **DNA marker 1** | 191Ir | SCIMAP core |
| **DNA marker 2** | 193Ir | SCIMAP core |

### Supplementary Table 6: Details of Statistical Analyses (Main Figures)

| Figure | Experiment | | Statistical test | p value |
|---|---|---|---|---|
| **1b** | BALB/c PPBC | Lung metastasis | t-test<br>WT=12 eIF4E$^{S209A}$=17 | p<0.0001 |
| **1c** | | Primary tumor out-growth | Two-way ANOVA<br>WT=12 eIF4E$^{S209A}$=17 | p=0.9992 |

| Figure | Experiment | | Statistical test | p value |
|---|---|---|---|---|
| **1d** | | Primary tumor Ki67 IHC | t-test WT=9 eIF4E$^{S209A}$=12 | p=0.5248 |
| **1e** | C57BL/6N PPBC | Lung metastasis | t-test WT=8 eIF4E$^{S209A}$=13 | p=0.8463 |
| **1f** | | Primary tumor out-growth | Two-way ANOVA WT=8 eIF4E$^{S209A}$=13 | p=0.0036 |
| **1g** | BALB/c PPBC (Endpoint) | Total T cells flow cytometry | t-test WT=5 eIF4E$^{S209A}$=8 | p=0.0036 |
| | | Cytotoxic T cells flow cytometry | t-test WT=5 eIF4E$^{S209A}$=8 | p=0.0202 |
| | | G-MDSCs flow cytometry | t-test WT=5 eIF4E$^{S209A}$=8 | p=0.0475 |
| | | M-MDSCs flow cytometry | t-test WT=5 eIF4E$^{S209A}$=8 | p=0.7557 |
| **2a** | BALB/c PPBC primary tumor (2 weeks) | IL-33 IHC | t-test WT=11 eIF4E$^{S209A}$=9 | WT versus eIF4E$^{S209A}$: p=0.0059 |
| **2b** | | Phospho-eIF4E/IL-33 IF | Spearman correlation WT=9 | r=0.9605 p<0.0001 |
| **2d** | Secreted IL33 in primary mammary gland fibroblasts | | n=2 | p=0.0133 |
| **2g** | 66cl4-Fibroblast co-culture MIA | | One-way ANOVA 3 independent experiments | 66cl4 alone versus WT: p<0.0001 WT versus eIF4E$^{S209A}$: p<0.0001 |
| **2h** | 66cl4-Fibroblast co-culture IL-33 qPCR | | Mann-Whitney test 3 independent experiments | p=0.0054 |
| **2k** | 66cl4-Fibroblast co-culture MIA | | t-test 3 independent experiments | p<0.0001 |

| Figure | Experiment | | Statistical test | p value |
|---|---|---|---|---|
| **2m** | MDA-MB-231-patient derived CAFs co-culture MIA | | t-test<br>1 invasion assay per donor. Every dot represents the cell numbers invaded per field | p<0.0001 for patients 1, 2, 3<br>p=00008 for patient 4 |
| **3a** | 66cl4-Fibroblast co-culture MIA<br>ST2 knockdown on 66cl4 | | One-way ANOVA | siCTL vs siIL1rl1-1: p=0.0403<br>siCTL vs siIL1rl1-2: p=0.0014 |
| **3c** | 66cl4 MIA -/+ rIL-33 | | Paired t-test<br>3 independent experiments | p=0.096 |
| **3d** | 66cl4 cells treated with rIL-33 | *Cxcl1* qPCR | Mann-Whitney test<br>3 independent experiments | p<0.0001 |
| | | *Ccl17* qPCR | | p<0.0001 |
| | | *Csf2* qPCR | | p<0.0001 |
| | | *Il6* qPCR | | p=0.0002 |
| | | *Il4* qPCR | | p=0.2249 |
| | | *Cxcl2* qPCR | | p=0.5406 |
| **3e** | Isolated WT MDSCs treated with rIL-33 for PD-L1 flow | | t-test<br>Vehicle Ctl=6<br>rIL-33 treated=6 | P=0.0260 |
| **3f** | ex vivo ILC2 activation | | Two-way ANOVA<br>2 independent experiments | IL13 secretion:<br>0μM vs 2.5μM SEL201: p=0.0012<br>0μM vs 5μM SEL201: p<0.001<br>2.5μM vs 2.5μM SEL201: p=0.0317<br>IL5 secretion:<br>0μM vs 5μM SEL201: p=0.0036 |
| **4a** | PPBC patient tumors | IF staining | Spearman correlation<br>9 PPBC patients | p4E/IL-33: r=0.8, p=0.0138 |
| **4b** | TCGA database | | Spearman correlation<br>1217 patients | r=0.2961, p<0.0001 |

| Figure | Experiment | | Statistical test | p value |
|---|---|---|---|---|
| **4c** | Human tumor microarray | IMC Phopho-eIF4E in tumor cells in BC, PrBC and PPBC | One-way ANOVA<br>BC cells=2393<br>PrBC cells=3579cells<br>PPBC cells=1206 cells | BC versus PrBC: p<0.0001<br>BC versus PPBC: p=0.0018<br>PrBC versus PPBC: p<0.0001 |
| | | IMC PD-L1 in tumor cells in BC, PrBC and PPBC | One-way ANOVA<br>BC cells=2393 cells<br>PrBC cells=3579cells<br>PPBC cells=1206 cells | BC versus PrBC: p<0.0001<br>BC versus PPBC: p=0.2947<br>PrBC versus PPBC: p<0.0001 |
| **4d** | Human tumor microarray | IMC CD8 proportion in BC, PrBC and PPBC | One-way ANOVA<br>BC patients=8<br>PrBC patients=6<br>PPBC patients=9 | BC versus PrBC: =0.3479<br>BC versus PPBC: p=0.0488<br>PrBC versus PPBC: p=0.3925 |
| | | IMC Phospho-eIF4E in CD8 cells in BC, PrBC and PPBC | One-way ANOVA<br>BC cells=535 cells<br>PrBC cells=794cells<br>PPBC cells=1911 cells | BC versus PrBC: p=0.2403<br>BC versus PPBC: p<0.0001<br>PrBC versus PPBC: p<0.0001 |
| **4e, f** | Human tumor microarray | IMC HLA-DR in CD8 cells in BC, PrBC and PPBC | One-way ANOVA<br>BC cells=535 cells<br>PrBC cells=794cells<br>PPBC cells=1911 cells | BC versus PrBC: =0.1703<br>BC versus PPBC: p<0.0001<br>PrBC versus PPBC: p<0.0001 |
| | | IMC PD-L1 in CD8 cells in BC, PrBC and PPBC | One-way ANOVA<br>BC cells=535 cells<br>PrBC cells=794cells<br>PPBC cells=1911 cells | BC versus PrBC: p=0.6594<br>BC versus PPBC: p<0.0001<br>PrBC versus PPBC: p<0.0001 |
| | | IMC correlation in CD8 cells between PD-L1 and HLA-DR | Spearman correlation<br>BC patients=8<br>PrBC patients=6<br>PPBC patients=9 | BC: r=0.2381, p=0.5821<br>PrBC: r= -0.0857, p=0.9194<br>PPBC: r=0.8, p=0.0138 |
| **5a** | BALB/c PPBC | Lung metastasis | One-way ANOVA<br>Control=10<br>Anti-PD-1=8<br>SEL201=12<br>Anti-PD-1+SEL201=10 | Control versus Anti-PD-1+SEL201: p=0.0074 |

| Figure | Experiment | | Statistical test | p value |
|---|---|---|---|---|
| **5b** | BALB/c PPBC: Lung metastasis phospho-eIF4E/CD8 IHC staining | CD8+ cell counts | One-way ANOVA<br>Control=7<br>Anti-PD-1=5<br>SEL201=7<br>Anti-PD-1+SEL201=7 | Control versus SEL201 p=0.0391<br>Control versus Anti-PD-1+SEL201 p=0.0642 |
| **5c** | BALB/c PPBC | Primary tumor outgrowth | One-way ANOVA<br>Control=10<br>Anti-PD-1=8<br>SEL201=12<br>Anti-PD-1+SEL201=10 | All comparisons are not significant |

**Supplementary Table 7: Details of Statistical Analyses (Supplementary Figures)**

| Figure ID | Experiment | | Statistical test | p value |
|---|---|---|---|---|
| **S.1a** | BALB/c PPBC | Lung metastasis | t-test<br>WT=12 eIF4E$^{S209A}$=17 | p<0.0001 |
| **S.1b** | | Tumor initiation rate | Mantel-Cox test<br>WT=10 eIF4E$^{S209A}$=17 | p=0.1013 |
| **S.1c** | | Primary tumor out-growth | t-test<br>WT=10 eIF4E$^{S209A}$=17 | p=0.4215 |
| **S.1e** | C57BL/6N PPBC | Primary tumor initiation | t-test<br>WT=10 eIF4E$^{S209A}$=17 | p=0.1445 |
| **S.1f** | | Primary tumor weight | t-test<br>WT=10 eIF4E$^{S209A}$=17 | P=0.7595 |
| **S.1f** | BALB/c involution | Adipocyte/Epithelium ratio | t-test per time points<br>WT=2 eIF4E$^{S209A}$=2 | ns |
| **S.3a** | E0771-Fibroblast co-culture MIA | | One-way ANOVA<br>3 independent experiments | E0771 alone versus WT: p<0.0375<br>WT versus eIF4E$^{S209A}$: p<0.0013 |
| **S.3b** | 66cl4-Fibroblast co-culture | | t-test<br>6 independent experiments | WT versus eIF4E$^{S209A}$ : p=0.0041 |
| **S.3e** | E0771-Fibroblast co-culture invasion | | t-test | p=0.0107 |
| **S.4a** | 66cl4-Fibroblast co-culture<br>ST2 knockdown qPCR | | One way anova<br>3 independent experiments | siCTL versus siST2-1: p=0.0059<br>siCTL versus siST2-2: p=0.0059 |
| **S.4c** | E0771 MIA -/+ rIL-33 | | t-test<br>3 independent experiments | p=0.0107 |
| **S.5b** | Human tumor microarray | IMC phospho-eIF4E in cell type proportion in BC and PrBC | One-way ANOVA<br>Tumor cells= 11703 (BC) and 4221 (PrBC) cells<br>Immune cells= 6455 (BC) and 9317 (PrBC) cells<br>Fibroblasts=149 (BC) and 61 (PrBC) cells<br>Pericytes=2843 (BC) and 2103 (PrBC) cells<br>Endothelial cell=8538 (BC) and 3922 (PrBC) cells | Tumor cells versus Immune cells CD45: p<0.0001 (BC) and p<0.0001 (PrBC)<br>Tumor cells versus Fibroblasts: p<0.0001 (BC) and p<0.0001 (PrBC)<br>Tumor cells versus Endothelial p<0.0001 (BC) and p<0.0001 (PrBC)<br>Tumor cells versus Other cells: p<0.0001 (BC) and p<0.0001 (PrBC) |

| Figure ID | Experiment | | Statistical test | p value |
|---|---|---|---|---|
| | | | Other cells=16130 (BC) and 6880 (PrBC) cells | |
| **S.5c** | Human tumor microarray | IMC eIF4E in cell type proportion in BC and PrBC and PPBC | One-way ANOVA<br>Tumor cells= 11703 (BC). 4221 (PrBC) and 7546 (PPBC) cells<br>Immune cells= 6455 (BC) , 9317 (PrBC) and 1179a (PPBC) cells<br>Fibroblasts=149 (BC), 61 (PrBC) and 44 (PPBC) cells<br>Pericytes=2843 (BC), 2103 (PrBC) and 2843 (PPBC) cells<br>Endothelial cell=8538 (BC), 3922 (PrBC) and 6665 (PPBC) cells<br>Other cells=16130 (BC), 6880 (PrBC) and 14246 (PPBC) cells | Tumor cells versus Immune cells CD45: p<0.0001 (BC), p<0.0001 (PrBC) and p<0.0001 (PPBC)<br>Tumor cells versus Fibroblasts: p<0.0313 (BC), p<0.0001 (PrBC) and p<0.0001 (PPBC)<br>Tumor cells versus Endothelial p<0.0001 (BC), p<0.0001 (PrBC) and p<0.0001 (PPBC)<br>Tumor cells versus Other cells: p<0.0001 (BC), p<0.0001 (PrBC) and p<0.0001 (PPBC) |
| **S.5d** | Human tumor microarray | IMC MNK1 in cell type proportion in BC and PrBC and PPBC | One-way ANOVA<br>Tumor cells= 11703 (BC). 4221 (PrBC) and 7546 (PPBC) cells<br>Immune cells= 6455 (BC) , 9317 (PrBC) and 1179a (PPBC) cells<br>Fibroblasts=149 (BC), 61 (PrBC) and 44 (PPBC) cells<br>Pericytes=2843 (BC), 2103 (PrBC) and 2843 (PPBC) cells<br>Endothelial cell=8538 (BC), 3922 (PrBC) and 6665 (PPBC) cells | Tumor cells versus Immune cells CD45: p<0.2506 (BC), p<0.0001 (PrBC) and p<0.0001 (PPBC)<br>Tumor cells versus Fibroblasts: p<0.0327 (BC), p<0.0001 (PrBC) and p<0.036 (PPBC)<br>Tumor cells versus Endothelial p<0.0001 (BC), p<0.0001 (PrBC) and p<0.0001 (PPBC)<br>Tumor cells versus Other cells: p<0.0001 (BC), p<0.0001 (PrBC) and p<0.0001 (PPBC) |

| Figure ID | Experiment | | Statistical test | p value |
|---|---|---|---|---|
| | | | Other cells=16130 (BC), 6880 (PrBC) and 14246 (PPBC) cells | |
| **S.5e** | Human tumor microarray | 99th percentile PD-L1 in tumor cells in BC, PrBC and PPBC | One-way ANOVA<br>BC=116 cells<br>PrBC =41cells<br>PPBC =75 cells | BC versus PrBC: p>0.9999<br>BC versus PPBC: p<0.0001<br>PrBC versus PPBC: p<0.0001 |
| **S.5g** | BALB/c PPBC: Lung metastasis phospho-eIF4E/CD8 IHC staining | phospho-eIF4E IHC score | One-way ANOVA<br>Control=7<br>Anti-PD1=5<br>SEL201=7<br>Anti-PD1+SEL201=7 | Control versus SEL201 p=0.0076<br>Control versus Anti-PD1+SEL201 p=0.0227 |
| **S.5h** | BALB/c PPBC | Primary tumor weight | One-way ANOVA<br>Control=10<br>Anti-PD1=8<br>SEL201=12<br>Anti-PD1+SEL201=10 | Control versus Anti-PD-1: p=0.9549<br>Control versus SEL201: p=0.6451<br>Control versus SEL201+Anti-PD-1: p=0.8256 |
