## Supplemental legends for "The pleiotropic effects of the MNK1/2-eIF4E axis support immune suppression and metastasis in a model of postpartum breast cancer"

**Supplementary Figure 1**

**a.** 66cl4 tumor cells are more metastatic when injected into the involuting mammary gland (i.e. the PPBC model) compared to the same cells injected into the virgin mammary gland. Data are presented as average tumor burden in the lung, each dot represents individual animals and the horizontal bar indicates the mean of the cohort. Two-tailed unpaired Student’s t test was used to calculate statistical significance (Scale bar=4mm, virgin n=9, PPBC n=12). **b.** Tumor free rates in WT and eIF4E$^{S209A}$ 66cl4 PPBC animals are similar. WT=12, eIF4E$^{S209A}$=16. **c.** 66cl4 primary tumor weights do not differ between WT and eIF4E$^{S209A}$ PPBC mice. WT=12, eIF4E$^{S209A}$=16. Data are presented as mean ± SD. **d.** E0771 cells implanted into the involuting mammary gland of p-eIF4E deficient PPBC hosts (C57BL/6N) give rise to fewer metastases than in WT PPBC animals. Two representative images of data presented in Figure 1e are shown. **e.** Tumor free rates in WT and eIF4E$^{S209A}$ E0771 PPBC animals WT=8, eIF4E$^{S209A}$=13. **f.** E0771 primary tumor weights do not differ between WT and eIF4E$^{S209A}$ PPBC mice. WT=8, eIF4E$^{S209A}$=13. Data are presented as mean ± SD. **g.** No difference in the ability of adipocytes to repopulate the mammary glands of WT and eIF4E$^{S209A}$ mice as they undergo involution. The average area of adipocytes and epithelial cells of 20 pictures for each time points were taken at a 10X magnification and analyzed using ImageJ software n=2 and data are presented as mean ± SD. **h.** WT and eIF4E$^{S209A}$ mammary glands do not differ in their gross morphology during involution. Images of the carmine red stained mammary gland are shown. **i.** The expression of p-STAT3 and STAT3 in mammary gland lysates was detected by western blot. Different times following the onset of involution is shown (LD8 = lactation day 8), ID = involution day 2, 4, 6. One representative experiment out of two independent experiments is shown.

**Supplementary figure 2**

Gating strategies used to define T cell and myeloid cell populations in the lungs of PPBC mice.

**Supplementary Figure 3**

**a.** E0771 cells invade less robustly in the presence of eIF4E$^{S209A}$ fibroblasts. Bars represent number of cells invaded per field of image and a summary of three biological replicates is shown. The horizontal bar indicates the mean. One-way ANOVA was used to calculate statistical significance. **b.** eIF4E$^{S209A}$ fibroblasts express less *Il33* compared to WT fibroblasts when co-cultured with E0771 cells. **c.** eIF4E$^{S209A}$ fibroblasts express less full-length and cleaved IL-33 compared to WT fibroblasts. IL-33 expression can be induced in fibroblasts upon co-culture with E0771 cells. One representative experiment of out three independent experiments is shown. **d.** SEL201 treatment reduces IL-33 protein levels in fibroblasts when co-cultured with E0771 cells. **e.** SEL201 treatment of WT fibroblasts suppresses E0771 invasion, when co-cultured. Bars represent the percentage of E0771 cells invaded compared to E0771 co-cultured with WT fibroblasts in absence of SEL201 (representative images are shown) and a summary of three biological replicates is shown. The horizontal bar indicates the mean. Student's t test was used to calculate statistical significance.

**Supplementary Figure 4**

**a.** IL-33 receptor ST2 (*Il1rl1*) knockdown in 66cl4 cells assessed by qRT-PCR. Data represent 3 biological replicates and are presented as mean mRNA fold change (normalized to housekeeping control *Rplp0*) ±SD. **b**. MNK1, p-ERK1/2, ERK1 and ERK2 protein levels in 66cl4 cells with or without rIL-33 treatment. One representative experiment out of three is shown. **c.** E0771 invasion is stimulated by rIL-33 treatment. Bars represent the percentage of E0771 cells invaded compared to E0771 cultured in absence of rIL-33 and a summary of three biological replicates is shown. Two-tailed unpaired Student's t test was used to calculate statistical significance.

**Supplementary Figure 5**

**a.** IMC staining showing an example of binding patterns of immune cell type markers: CD68 (red) and CD3 (green), which are mutually exclusive. CD8a (green) and CD4 (blue), which are mutually exclusive, and CD4 (blue) and FoxP3 (red), which co-occur. Proportion of different cell types and their level of expression of phospho-eIF4E **(b)**, eIF4E **(c)**, and MNK1 **(d)** in BC (8 patients), PrBC (6 patients), PPBC (9 patients). Each dot represents one cell. One-way ANOVA was used to calculate statistical significance. **e.** PD-L1 expression in the $99^{th}$ percentile of phospho-eIF4E-expressing tumor cells. Each dot represents one cell. One-way ANOVA was used to calculate statistical significance. **f.** Schematic of the therapeutic regimen (anti-PD-1 and SEL201 administration) used in PPBC mice. **g.** SEL201 treatment efficiently repressed phospho-eIF4E expression. Each dot in the graphs represents one animal and the horizontal bar indicates the mean of the cohort. One-way ANOVA was used to calculate statistical significance. Vehicle group n=7, anti-PD-1 treated group n=5, SEL201-treated group n=7, SEL201+anti-PD-1 group n=7. **h**. SEL201 and/or anti-PD-1 treatment did not alter the primary 66cl4 tumor weight. Data are presented as mean ±SD. Vehicle group n=10, anti-PD-1 treated group n=8, SEL201-treated group n=12, SEL201+anti-PD-1 group n=10. **i**. Treatment with SEL201 and/or anti-PD-1 blockade does not significantly affect body weight. Data shown represent average body weight of animals in each group and is presented as mean ±SEM.
