## Supplemental figures for "The pleiotropic effects of the MNK1/2-eIF4E axis support immune suppression and metastasis in a model of postpartum breast cancer"

Supplementary Figure 1

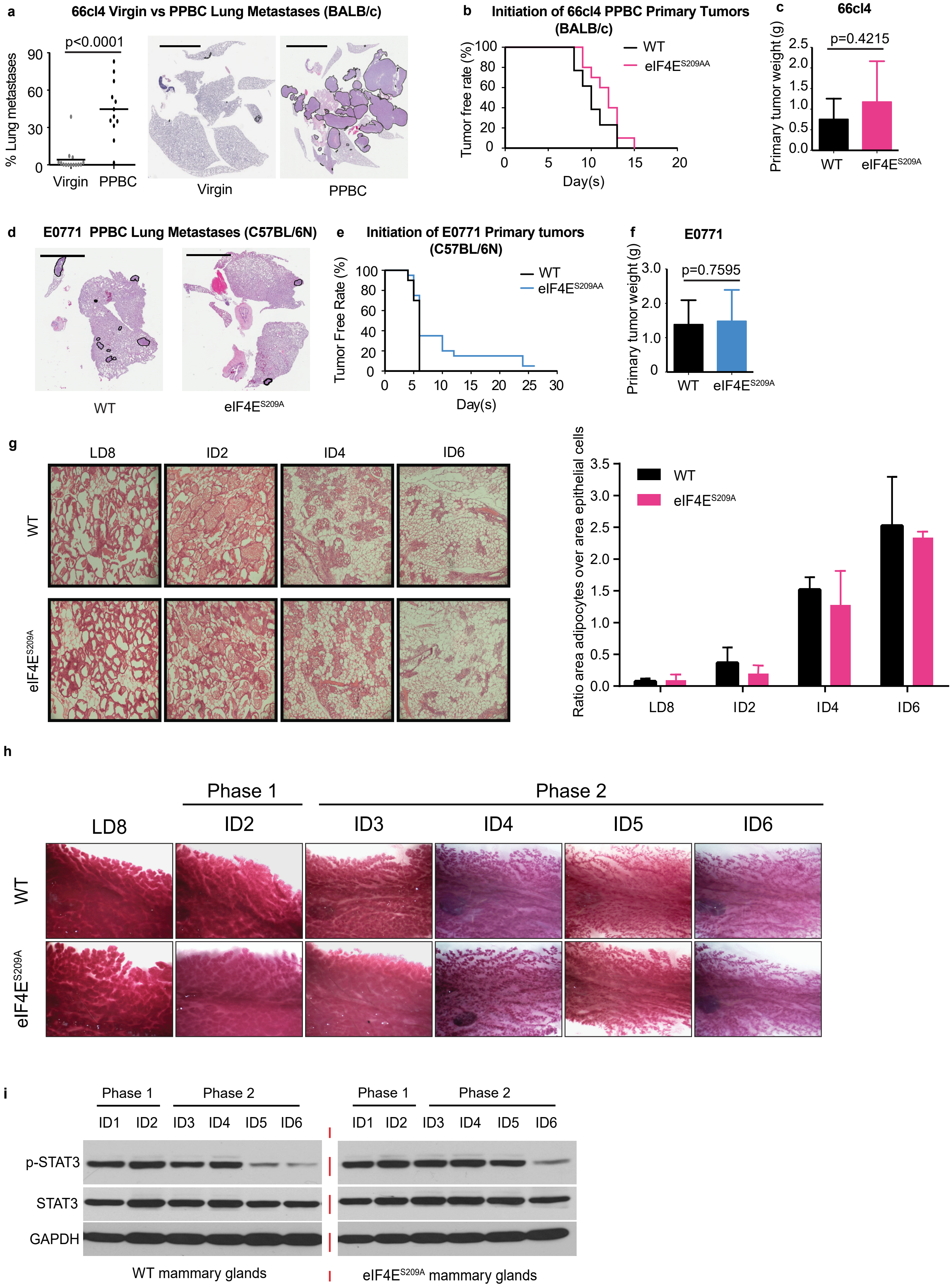

**Supplementary Figure 2**

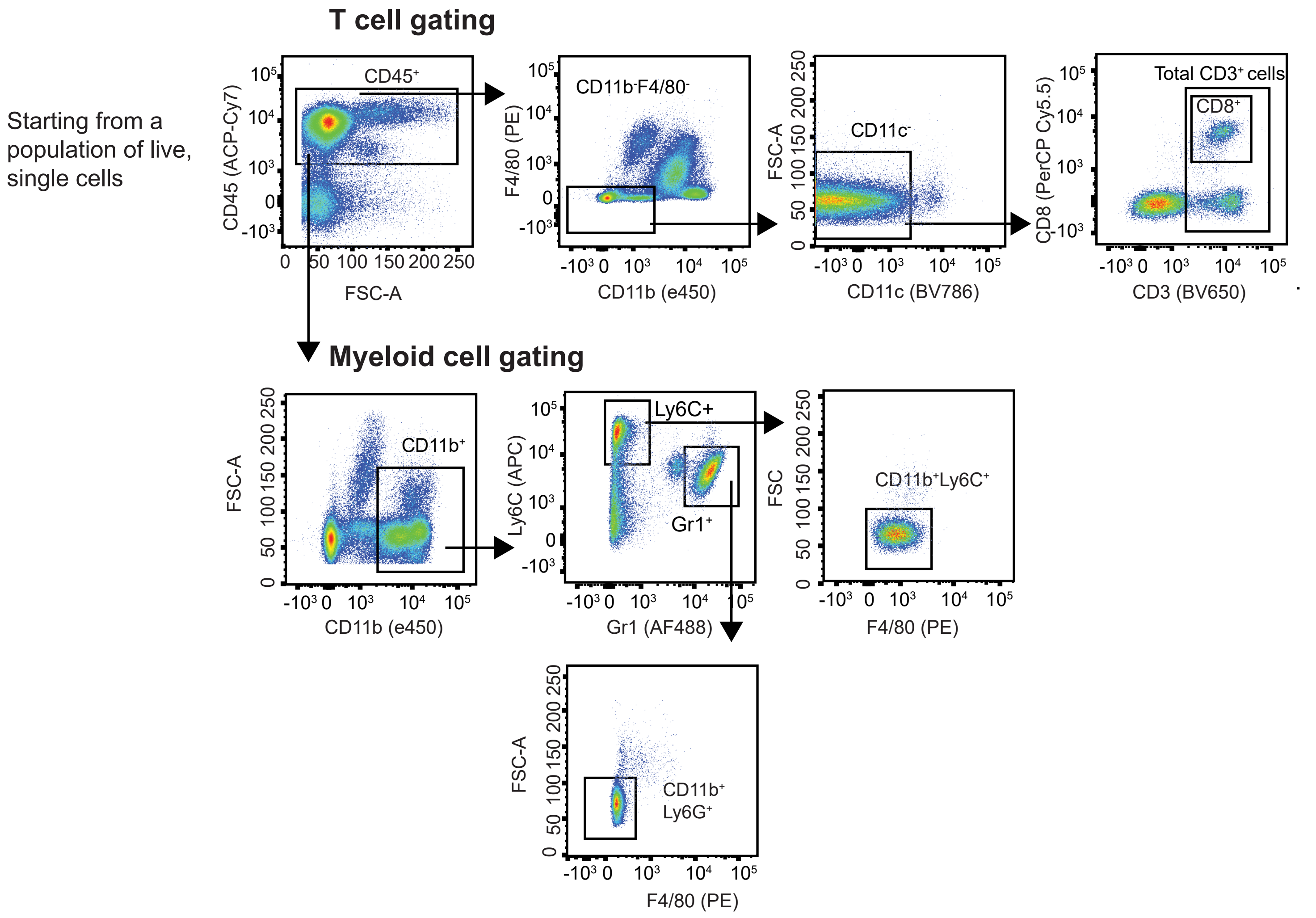

**Supplementary Figure 3**

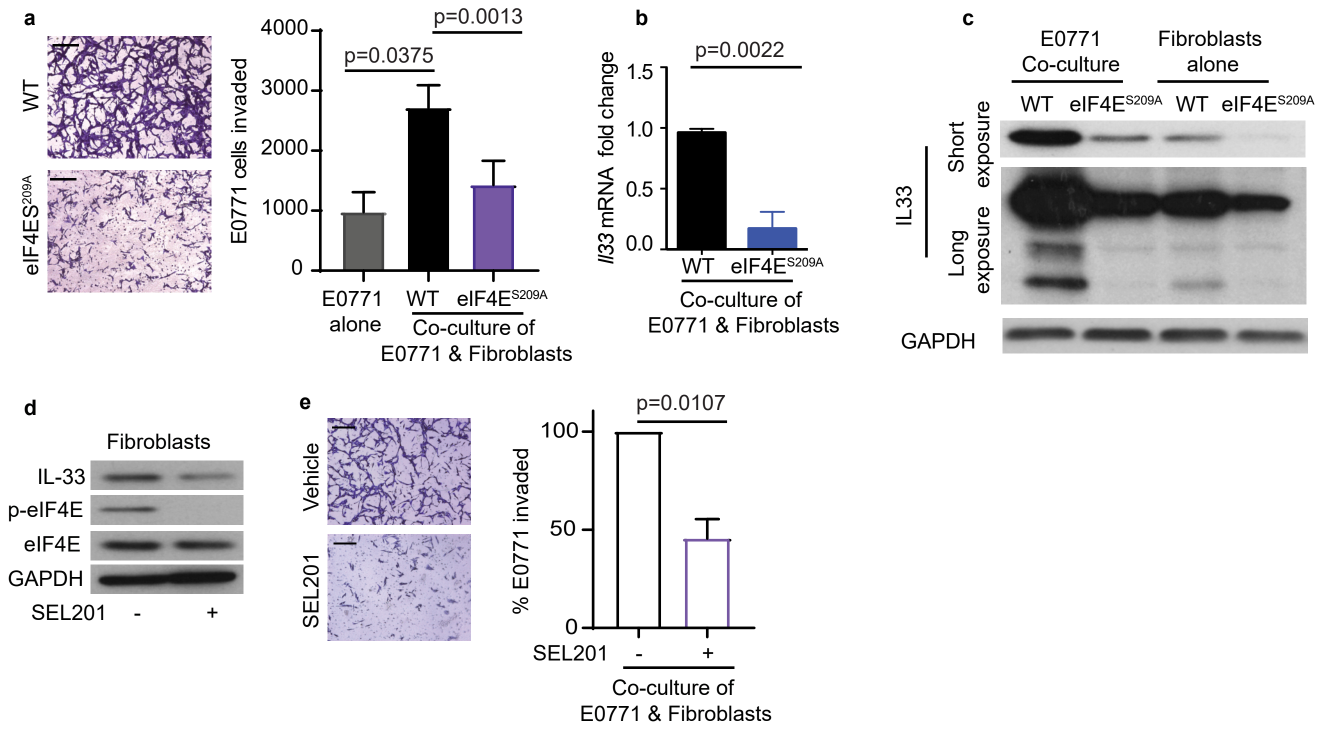

**Supplementary Figure 4**

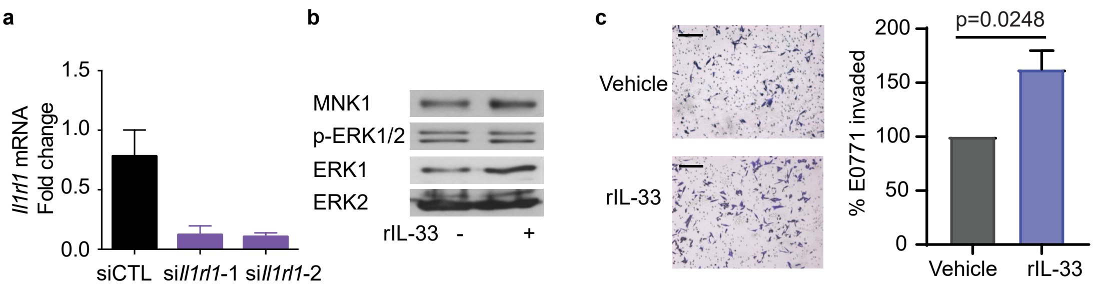

**Supplementary Figure 5**

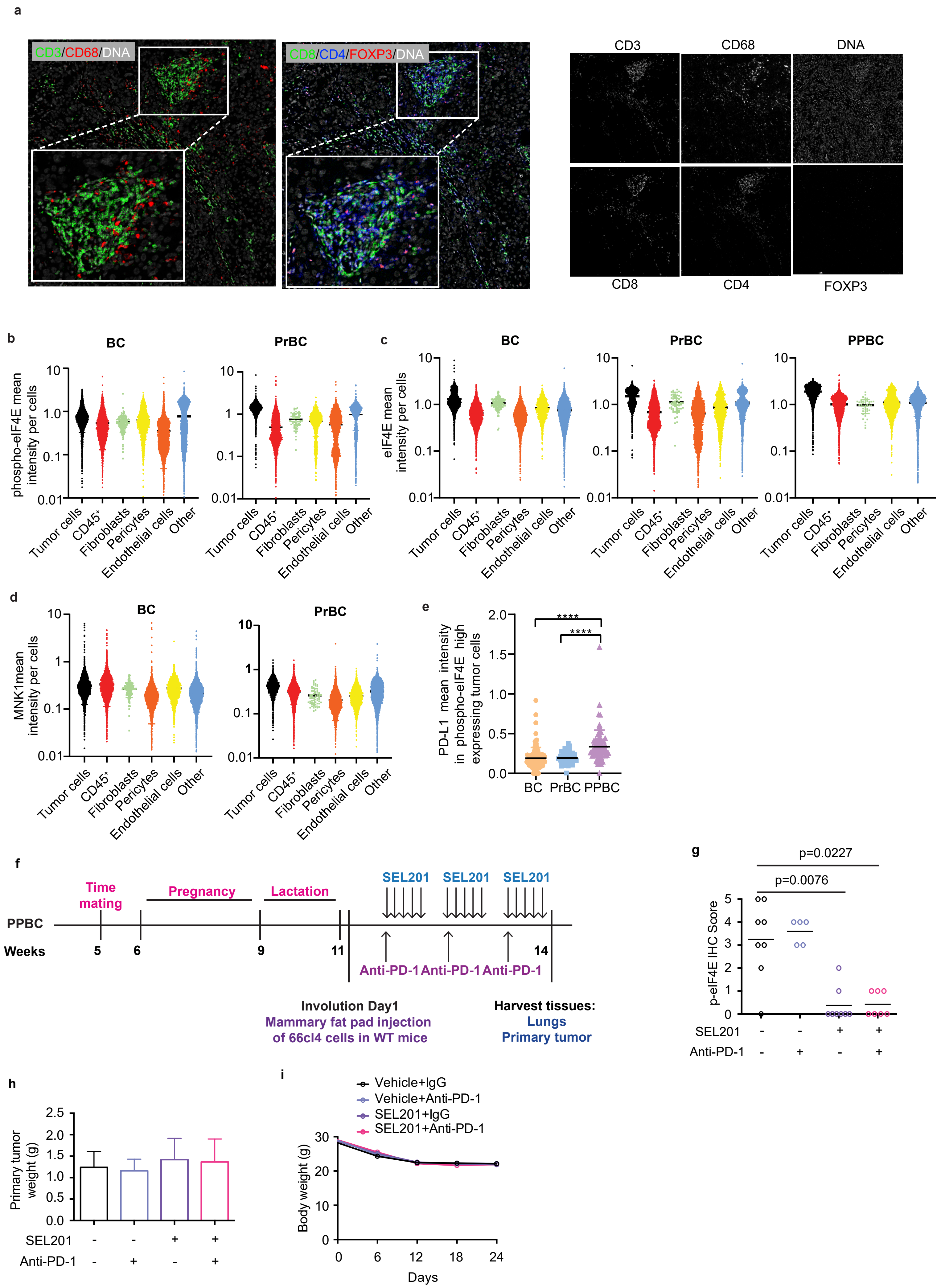
